# Network-level encoding of local neurotransmitters in cortical astrocytes

**DOI:** 10.1101/2023.12.01.568932

**Authors:** Michelle K. Cahill, Max Collard, Vincent Tse, Michael E. Reitman, Roberto Etchenique, Christoph Kirst, Kira E. Poskanzer

## Abstract

Astrocytes—the most abundant non-neuronal cell type in the mammalian brain—are crucial circuit components that respond to and modulate neuronal activity via calcium (Ca^2+^) signaling^1–8^. Astrocyte Ca^2+^ activity is highly heterogeneous and occurs across multiple spatiotemporal scales: from fast, subcellular activity^3,4^ to slow, synchronized activity that travels across connected astrocyte networks^9–11^. Furthermore, astrocyte network activity has been shown to influence a wide range of processes^5,8,12^. While astrocyte network activity has important implications for neuronal circuit function, the inputs that drive astrocyte network dynamics remain unclear. Here we used *ex vivo* and *in vivo* two-photon Ca^2+^ imaging of astrocytes while mimicking neuronal neurotransmitter inputs at multiple spatiotemporal scales. We find that brief, subcellular inputs of GABA and glutamate lead to widespread, long-lasting astrocyte Ca^2+^ responses beyond an individual stimulated cell. Further, we find that a key subset of Ca^2+^ activity—propagative events—differentiates astrocyte network responses to these two major neurotransmitters, and gates responses to future inputs. Together, our results demonstrate that local, transient neurotransmitter inputs are encoded by broad cortical astrocyte networks over the course of minutes, contributing to accumulating evidence across multiple model organisms that significant astrocyte-neuron communication occurs across slow, network-level spatiotemporal scales^13–15^. We anticipate that this study will be a starting point for future studies investigating the link between specific astrocyte Ca^2+^ activity and specific astrocyte functional outputs, which could build a consistent framework for astrocytic modulation of neuronal activity.

## Main Text

A set of defined rules governing neuronal input-output relationships is a cornerstone of neuroscience. Given a specific excitatory or inhibitory neurotransmitter (NT) input, post-synaptic membrane potential changes that lead to action potentials can be accurately predicted. But, neurons are not the only cells in the nervous system that sense NTs. Astrocytes—the most abundant non-neuronal cell type in the mammalian brain—are crucial circuit components that respond to and modulate neuronal activity via calcium (Ca^2+^) signaling^1–8^. However, the set of rules governing input-output relationships in astrocytes is poorly defined, in part because it’s unclear over which spatiotemporal scales these relationships should be evaluated. While there appear to be fast and local astrocytic responses to local stimuli^3,4^, there is also evidence to suggest that astrocyte responses to local stimuli have a spatiotemporally distributed component, as local astrocyte stimulation can lead to distributed changes in neuronal activity and plasticity^16,17^. Thus, a comprehensive framework describing input-output relationships in astrocytes requires simultaneous investigation of activity across multiple spatiotemporal scales.

Here, we set out to build an input framework governing transient and sustained cortical astrocyte Ca^2+^ activity at three spatial scales: subcellular, single cell, and network. To take a physiologically relevant and comparative approach, we focused on astrocyte responses to the two major NTs: glutamate and GABA. While previous studies demonstrate general astrocyte Ca^2+^ increases in response to these NTs^2,6,18,19^, our goal was to link specific excitatory and inhibitory chemical inputs to specific astrocyte Ca^2+^ activity, and map the scales over which astrocytes could exert effects on neuronal circuitry.

### NTs drive distinct astrocyte Ca^2+^ activity

To first test whether astrocytes show generally distinct activity in response to different NTs, we used two-photon (2P) Ca^2+^ imaging (via the genetically encoded intracellular indicator, Cyto-GCaMP6f) of astrocytes while sequentially bath-applying GABA and glutamate receptor agonists onto *ex vivo* acute cortical slices from mice (Fig. 1a). We activated the GABAergic and glutamatergic GPCRs expressed by astrocytes^20,21^ (Extended Data Fig. 1a, Supplementary Videos 1 and 2), using baclofen to activate GABA_B_ receptors^2,18,22^ and a broad spectrum metabotropic glutamate receptor (mGluR) agonist, (1S-3R)-ACPD (t-ACPD)^16,23–25^, to activate mGluR_3_, the mGluR subtype expressed by astrocytes at this age^26^, while silencing neuronal firing with tetrodotoxin (TTX). We analyzed the resulting Ca^2+^ activity using the event-detection software AQuA^9^ (Fig. 1b). In the same populations of astrocytes, with similar levels of baseline activity (Extended Data Fig. 1b), GABA_B_R or mGluR_3_ activation both increased Ca^2+^ event frequency, but each led to distinct Ca^2+^ responses in time-course and magnitude. Using both event-based and region-of-interest (ROI)-based analysis methods, we found with t-ACPD, Ca^2+^ activity increases were robust and transient, whereas baclofen caused a delayed and prolonged activation, lasting through the end of recording (Fig. 1c, Extended Data Fig. 1c–e). Analyzing individual Ca^2+^ events by area and duration, we found a population of events larger and longer compared to baseline with t-ACPD, but not with baclofen (Fig. 1d, Extended Data Fig. 1f,g). To ensure that these distinct responses weren’t dependent on a specific agonist concentration or agonist order, we quantified activity changes across a broad concentration range, alternating agonist order between concentrations. Across Ca^2+^ event features, we saw a consistently higher response with mGluR_3_ compared to GABA_B_R activation (Fig. 1e–h), demonstrating that the same cortical astrocyte populations exhibit distinct activity, with distinct time courses, in response to different NTs.

**Fig. 1.**
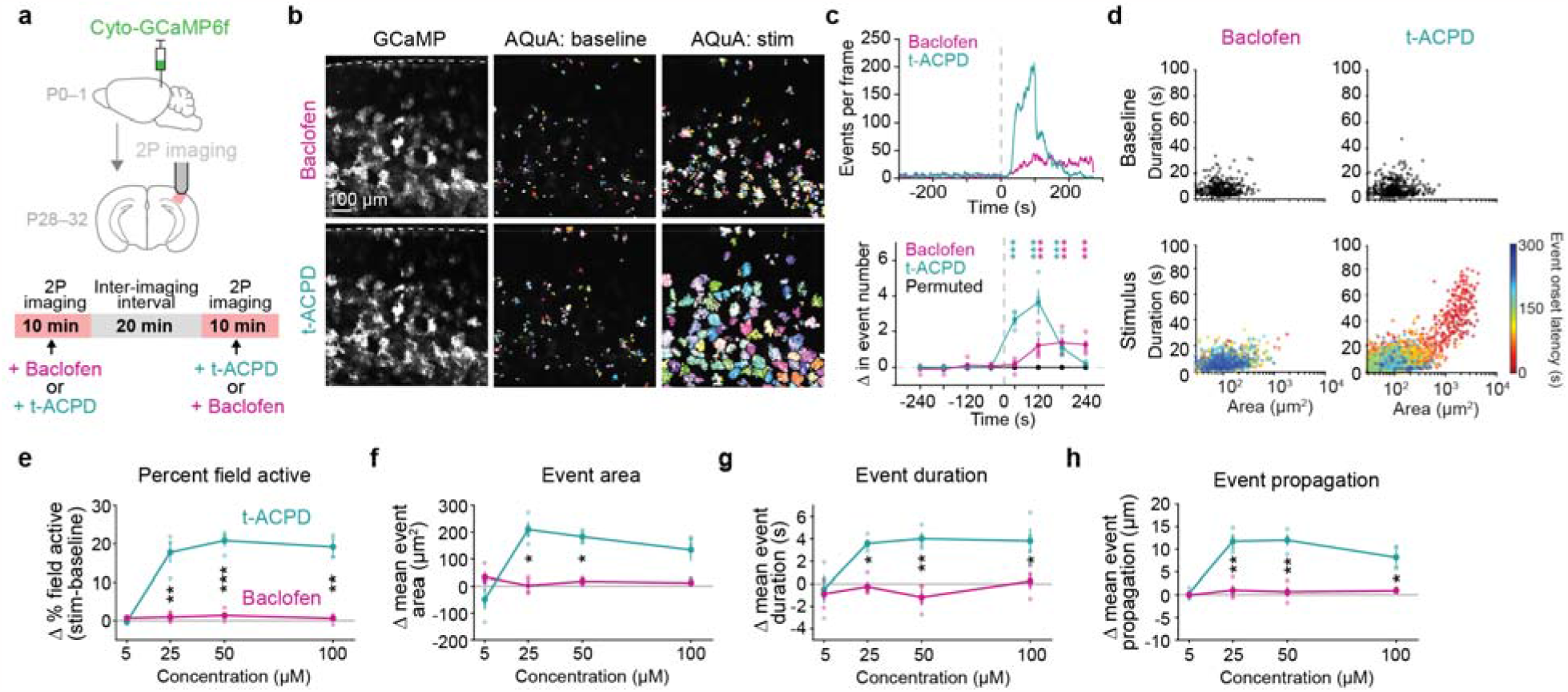
Direct GABAergic and glutamatergic receptor activation drive distinct astrocyte Ca^2+^ activity. **a**, Experimental strategy for expression of cyto-GCaMP6f and 2P imaging astrocytic Ca^2+^ in acute V1 cortical slices during pharmacological activation via bath-application. Receptor agonists were sequentially bath-applied to the same slice, with an inter-imaging interval of >20min, including >10min wash-out period. **b**, Representative astrocytic GCaMP6f fluorescence (left column) during bath-application of GABA_B_-specific agonist baclofen (50μM, top row) and mGluR agonist t-ACPD (50μM, bottom row). Dotted line denotes pia. All AQuA-detected events 300s before (middle column) and after (right column) bath-application of agonists, from the same slice shown on left. **c**, Top: Representative time-series traces (AQuA events per frame) of FOVs in B. Bottom: Average change in the events/minute compared to average baseline. 300–0s before and 0–240s after bath-application of agonists used to calculate change in events/60s for each active astrocyte (≥ 1 AQuA-detected event) and averaged for each slice. Data shown by slice (n = 4 slices stimulated with 50μM agonist, light dots) and mean ± standard error of the mean (sem) (solid dots and error bars). Permutation test used to determine significance. *p*-values for all timepoints are in Extended Data Table 1. All traces are aligned to 0s, the frame of agonist entry in the imaging chamber. **d**, Scatter plots of the area and duration of individual Ca^2+^ events at baseline (top row, black) and after bath-application of baclofen (bottom left) or t-ACPD (bottom right). Bottom row: events following bath-application of agonists color-coded by onset time. Dots represent individual Ca^2+^ events from *n* = 4 slices stimulated with 50μM agonist. **e–h**, Dose-response curves showing average change in Ca^2+^ features with bath-application of baclofen (pink) or t-ACPD (green) at four concentrations. Agonist order alternated between conditions: Baclofen was added first at 5 and 50μM and second at 25 and 100μM. Change calculated by comparing 120s before and after agonist entry. Data shown by slice (*n* = 4 slices, 4 mice for each concentration, light dots) and mean ± sem (solid dots and error bars). Paired t-tests at each concentration compare activity changes induced by each agonist. *p*-values corrected for multiple comparisons using Bonferroni-Holm correction with FWER ≤ 0.05. *p*-values for all concentrations and features are in Extended Data Table 2.

GABA_B_R and mGluR_3_ are both G_i_-coupled GPCRs canonically linked to decreases in intracellular cAMP. To explore whether these two G_i_-GPCRs also engage cAMP in NT-specific ways, we expressed the genetically encoded cAMP sensor Pink Flamindo^27^ in astrocytes, and bath-applied agonists selective for these receptors (Extended Data Fig. 1h–k). We switched from using a broad spectrum mGluR agonist, t-ACPD (Fig. 1), to an mGluR_3_-selective agonist, LY 379268 (Extended Data Fig. 1h–k), to specifically examine the effect of this G_i_-GPCR activation on cAMP activity. In contrast to canonical G_i_-GPCR signaling, we saw slow and sustained cAMP increases^28,29^ with both agonists, with more cells responding to mGluR_3_ compared to GABA_B_R activation (Extended Data Fig. 1j). When comparing astrocytic Ca^2+^ and cAMP signaling in response to agonists, we found significantly more dynamic Ca^2+^ activity compared to cAMP (Extended Data Fig. 1k). Although Ca^2+^ isn’t a canonical downstream signaling partner of G_i_-GPCRs, our results confirm previous findings that astrocytes do signal via mGluR_3_ and GABA_B_R to mobilize intracellular Ca^2+^ (ref. ^2,18,28,30^) potentially via PLC signaling^31,32^ or by βγ subunits directly binding to IP_3_R^33,34^. The relative lack of dynamism in cAMP compared to Ca^2+^ led us to focus only on Ca^2+^ as the second messenger more likely to exhibit NT-specific responses to spatiotemporally restricted—and more physiological—NT release.

### Single astrocytes respond to subcellular release of NTs

To release NTs with spatiotemporal precision, we used 2P photo-release (“uncaging”) of caged neurotransmitters (Fig. 2a), as is commonly used to interrogate post-synaptic physiology via restricted activation area and duration^7,16,35–39^. To compare the effects of GABA or glutamate on the same astrocytes, we chose a class of caged compounds (with ruthenium bipyridine [RuBi] backbones), bound to either GABA^40,41^ or glutamate^41,42^, that can be 2P-uncaged (800nm) during simultaneous GCaMP Ca^2+^ imaging with a second 2P laser (excitation 980nm) (Fig. 2b). With this strategy, the uncaging/imaging experimental paradigm is common to both GABA and glutamate conditions. Because of likely variability in the Ca^2+^ response to NT across individual cells^7,43^, we imaged the same astrocytes while sequentially uncaging GABA and glutamate at the same subcellular location, separated by an inter-imaging interval of > 20 min, including a washout period of > 10 min. To account for any changes resulting from prior NT release, we alternated the order of GABA- or glutamate-uncaging between slices. To first quantify the properties of NT release in this dual-2P uncaging/imaging strategy, we imaged an extracellular-facing glutamate sensor (GluSnFR^44^) while uncaging RuBi-glutamate (Fig. 2c). We confirmed that NT release was spatiotemporally confined at the intended location over an area of ∼25 μm^2^ and duration of 0.5–1 s (Fig. 2d). To ensure that the uncaging laser itself does not stimulate astrocytes, we also stimulated GCaMP-expressing astrocytes with the uncaging laser alone in the absence of RuBis, and did not observe a change in average Ca^2+^ fluorescence or event frequency (Extended Data Fig. 2a,b).

**Fig. 2.**
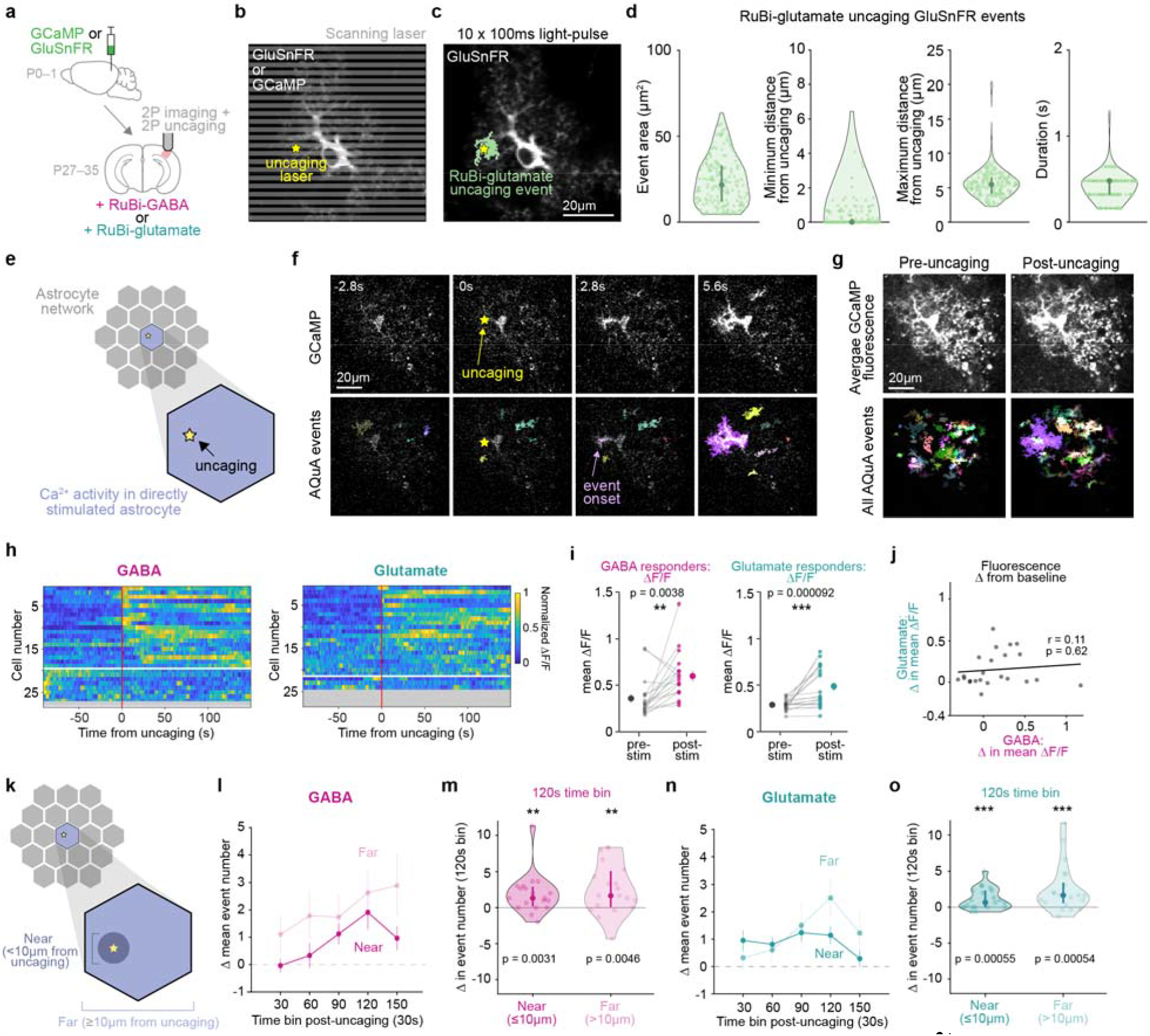
Subcellular, spatiotemporally restricted release of NTs increases Ca^2+^ activity within directly stimulated astrocytes. **a**, Experimental strategy for simultaneous 2P imaging of astrocyte Ca^2+^ (Cyto-GCaMP6f) or extracellular glutamate (GluSnFR), and 2P uncaging of NTs in acute V1 cortical slices. **b**, Imaging/uncaging schematic. Gray lines = scanning laser. Yellow star = NT uncaging site. **c**, To validate spatial precision of 2P uncaging, RuBi-glutamate was uncaged at a GluSnFR-expressing astrocyte. Yellow star = uncaging laser location; green = GluSnFR event footprint post-uncaging. **d**, GluSnFR event features after RuBi-glutamate uncaging. Data shown by individual glutamate events, median, 25^th^ and 75^th^ percentile (*n* = 72 trials, 12 recordings, 4 slices, 2 mice). **e**, Schematic illustrating analysis throughout figure is of Ca^2+^ activity from astrocyte directly stimulated by uncaging. **f**, Representative GCaMP6f fluorescence time-course in an individual astrocyte seconds before and after RuBi-GABA uncaging. Top row: raw fluorescence; bottom row: overlaid AQuA events. Yellow star = uncaging location and frame. **g**, Average GCaMP fluorescence 150–0s pre-uncaging (left column) and 0–150s post-uncaging (right column) from astrocyte in **f**. Top row: raw fluorescence; bottom row: all AQuA-detected Ca^2+^ events. **h**, Ca^2+^ activity in astrocytes directly stimulated by GABA (left) or glutamate (right) uncaging. Each row shows average ΔF/F traces from AQuA-detected events/cell, normalized individually between 0–1 for each cell. Cells sorted by onset time (first post-stim peak ≥ threshold [mean baseline ΔF/F + 3SD], with threshold calculated by cell). Red line = NT uncaging time. White line separates responding (above) and non-responding cells (below). Responders are defined as any astrocyte with ≥ 1 post-stim frame with ∆F/F ≥ threshold. Greyed-out rows represent cells excluded due to significant event frequency increases or decreases during the baseline period (see *2P uncaging event-based analysis* methods). **i**, Mean fluorescence pre-and post-stim from astrocytes responding to direct GABA (left) or glutamate (right) uncaging. 90–0s before and 0–150s after uncaging used to calculate mean ∆F/F pre- and post-stim/astrocyte. Data shown by cell (light dots and grey lines) and mean ± sem (dark dots and error bars). (For **i, l–o**, *n* = 19/27 directly stimulated cells responded to GABA and 21/24 to glutamate from *n* = 7 slices, 4 mice). Wilcoxon signed-rank test compare pre-and post-stim values. **j**, Fluorescence change in directly stimulated astrocytes following GABA and glutamate uncaging. 90–0s before and 0–150s after uncaging used to calculate mean change per cell. Pearson’s correlation shows no significant relationship between fluorescence change following GABA and glutamate uncaging (*p* = 0.62). **k**, Schematic illustrating that Ca^2+^ events within directly stimulated astrocyte are divided into events *near* and *far* from uncaging site. Note: events far from uncaging site are outside the radius of NT spread (d, “maximum distance from uncaging”), but still within the directly stimulated cell. **l, n**, Event frequency change near and far from GABA (**l**) and glutamate (**n**) uncaging within responding, directly stimulated cells. 90–0s before and 0–150s after uncaging used to calculate event number/30s. Data shown as mean ± sem. **m, o**, Event frequency change during a period of generally high activity (90–120s after uncaging, “120s” bin) from **l & n**. Data shown by cell, median, 25^th^ and 75^th^ percentile. Wilcoxon signed-rank test compare change from baseline for each cell compartment.

After validating the spatiotemporal precision of this approach, we next released NT during GCaMP imaging. We first analyzed the Ca^2+^ activity within the astrocyte that was directly stimulated (Fig. 2e). We observed examples of Ca^2+^ increases within seconds, in close proximity to the uncaging site (Fig. 2f,g and Supplementary Videos 3 and 4). By plotting ∆F/F and sorting by latency-to-fluorescence increases, we saw most astrocytes increase Ca^2+^ activity following NT release (Fig. 2h, above white line [70% and 88% of cells for GABA and glutamate, respectively], 2i), but the area and duration of Ca^2+^ events were unchanged (Extended Data Fig. 2e). The activity increases often lasted for 2.5 minutes after NT release, the post-uncaging duration of the recording (Fig. 2h, Extended Data Fig. 2b), validating previous findings that NT-induced astrocyte Ca^2+^ activity can be long-lasting^2,6^. Comparing the same astrocyte’s response to both NTs, we found no significant relationship between the magnitude of its response to GABA vs. glutamate (Fig. 2j), a controlled comparison given similar levels of activity within each cell prior to uncaging (Extended Data Fig. 2c,d). To confirm that the Ca^2+^ elevations were due to activation of astrocytic GPCRs, we next performed NT uncaging in slices where GABA_B_R or mGluR were inhibited pharmacologically, and found that Ca^2+^ increases were indeed blocked in these conditions (Extended Data Fig. 2a,b).

Astrocyte Ca^2+^ activity can be highly compartmentalized^3,4,7,43^, so we next tested whether observed changes in Ca^2+^ activity within the stimulated astrocyte were confined to subcellular regions directly exposed to initial NT release (<10μm from uncaging, Fig. 2c,d). We found an increased frequency of Ca^2+^ events both near (<10μm) and far (≥10μm) from the uncaging site (Fig. 2k–o and Extended Data Fig. 2f), with increases in both spatial domains peaking ≥ 1 min after uncaging for both NTs. These data demonstrate that spatiotemporally restricted NT release can drive Ca^2+^ activity in subcellular compartments extending beyond the stimulated region.

### Astrocyte networks respond to subcellular NTs

To ask whether activity changes extended beyond single cells, we next investigated population-wide Ca^2+^ activity in neighboring astrocytes within the gap junctionally coupled local network (Fig. 3a). Within the 300×300μm imaging field-of-view (FOV), the astrocyte over which NT was uncaged was approximately centered. Neighboring astrocytes (n=10.3±3.85, mean±SD) with GCaMP6f activity were imaged and distinguished from the uncaged cell by delineating cell maps. The active neighboring astrocytes within a given FOV define a “local network” (Fig. 3a,b). We observed general Ca^2+^ increases within the local network of astrocytes after uncaging (Fig. 3b, Extended Data Fig. 3d–f and Supplementary Videos 5 and 6). While we saw heterogeneity in the timing and magnitude of local network responses to subcellular NT release in the uncaged cell, the majority of imaged networks responded with population-wide fluorescence increases (Fig. 3c, left).

**Fig. 3.**
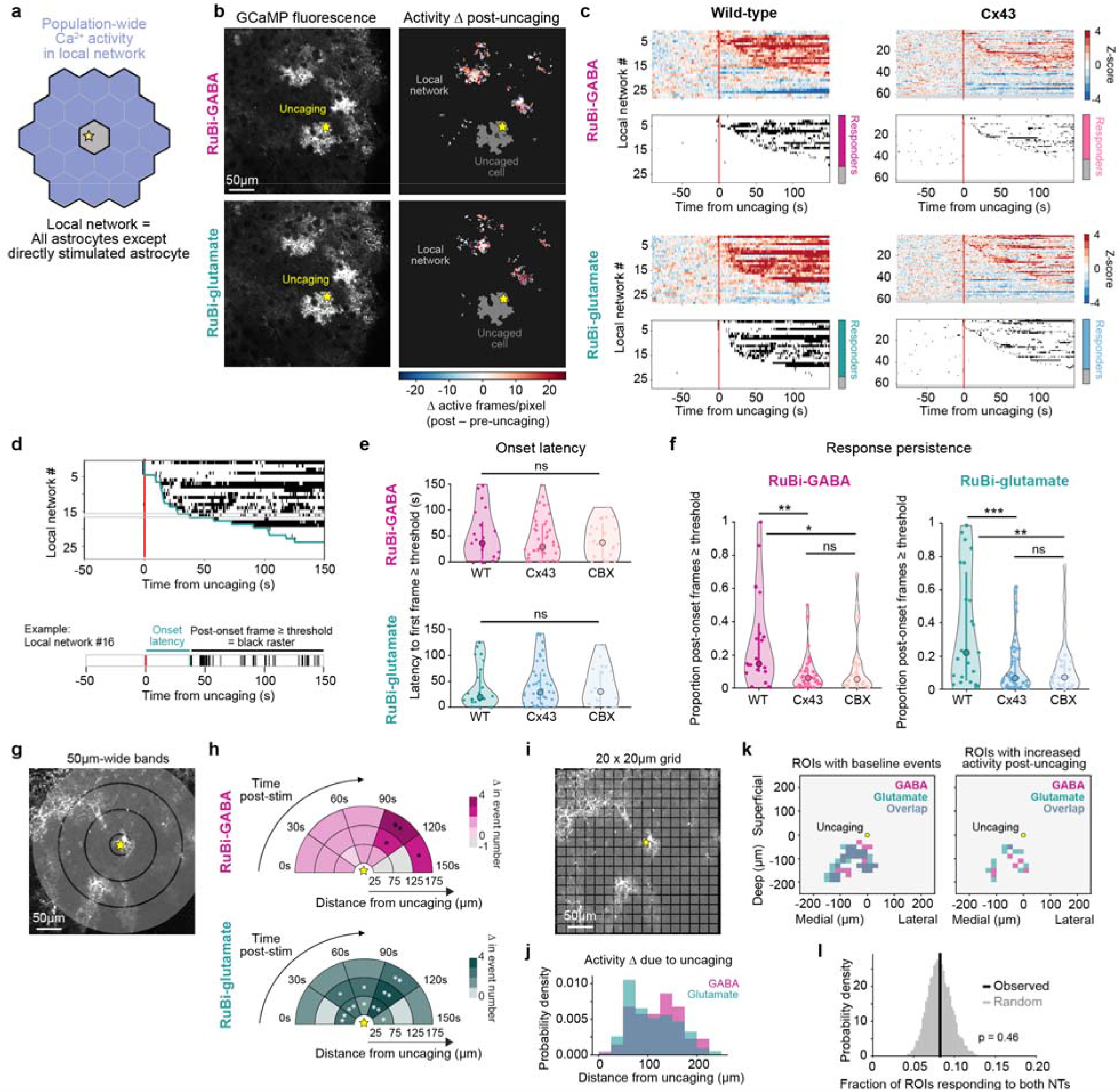
Subcellular release of GABA and glutamate increases Ca^2+^ activity in the local astrocyte network via Cx43. **a**, Schematic illustrating analysis throughout figure is of population-wide Ca^2+^ activity of all astrocytes in the FOV that are not directly stimulated by uncaging (*local network*). **b**, Left: Representative astrocytic GCaMP6f fluorescence in a V1 slice during 2P uncaging of RuBi-GABA (top) and RuBi-glutamate (bottom). Right: Representative spatial heatmaps of Ca^2+^ activity changes in the local astrocyte network from left following RuBi-GABA (top) and RuBi-glutamate (bottom) uncaging. Yellow star = uncaging site, same for each NT. Colors denote change in active frame number/pixel (all AQuA-detected events 150s before and after uncaging; red = activity increase, blue = decrease; activity in the uncaged cell [dark grey] is excluded). **c**, Top: Ca^2+^ activity from all recorded local astrocyte networks. Each row shows activity of one local network as the z-score of the mean ΔF/F from AQuA-detected events in the network. Mean ∆F/F and SD calculated using a baseline period 90–0s before uncaging. Networks sorted by onset time (first post-stim peak ≥ threshold [mean baseline ΔF/F + 3 SD]), with threshold calculated by trial). Red line = NT uncaging time. Greyed-out rows represent networks in which no events were detected outside of the uncaged cell. Bottom: binarized raster plots show each frame ΔF/F ≥ threshold. Stacked bar graphs show proportion of local networks exhibiting an initial fluorescence increase following uncaging (responder). Responders defined as any network with ≥1 post-stim frame ∆F/F ≥ threshold. WT (left) and Cx43^floxed^ slices (right). Fisher’s exact test compares proportion of responders across conditions: *p* = 0.62 (GABA WT vs Cx43^floxed^), *p* = 0.78 (glutamate WT vs Cx43^floxed^), *p* = 0.75 (GABA WT vs glutamate WT). **d**, Top: example binarized raster plot from C. Green line = response onset for each network (first post-stim frame ≥ threshold). Bottom: Example local network, showing onset latency (green) as time between NT-uncaging and response onset, and post-onset frames ≥ threshold (black tick marks). **e**, One-way ANOVA compares the onset latency across conditions (WT, Cx43^floxed^, and CBX). *p* = 0.82 (GABA), *p* = 0.89 (glutamate). (For **e, f**, Data shown by responding network, median, and 25^th^ and 75^th^ percentile. WT: *n* = 21 networks responding to GABA and 23 networks to glutamate; *n* = 7 slices, 4 mice; Cx43^floxed^: *n* = 42 networks responding to GABA and 47 networks to glutamate; *n* = 16 slices, 8 mice; CBX: *n* = 24 networks responding to GABA and 24 networks to glutamate; *n* = 8 slices, 4 mice). **f**, Persistence of network-level responses calculated as the proportion of post-onset frames ≥ threshold. One-way ANOVA followed by Tukey-Kramer Test determine significant pairwise comparisons between conditions for each NT. GABA: *p* = 0.0010 (WT v Cx43^floxed^), 0.025 (WT v CBX) and 0.72 (Cx43^floxed^ v CBX). Glutamate: *p* = 0.00034 (WT v Cx43^floxed^), 0.0032 (WT v CBX) and 0.98 (Cx43^floxed^ v CBX). **g**, Sholl-like analysis schematic. Grey concentric circles = 50μm bands. Yellow star = NT uncaging site. Inner radius of innermost band begins 25μm from uncaging, as events <25μm from the uncaging site are likely to occur within stimulated astrocyte. Outer radius of outermost band is 175μm from uncaging, as >175μm from the uncaging site can be outside the FOV; see Extended Data Fig. 3i. **h**, Ca^2+^ event frequency change in the local network after GABA (top) and glutamate (bottom) uncaging. 90–0s before and 0–150s after uncaging used to calculate event number/30s. Permutation test used to determine significance. *p*-values for all timepoints and bands are in Extended Data Table 6. **i**, Grid-based ROI (20 x 20 μm) schematic. **j**, Distribution of distances from uncaging site to center of ROIs active post-uncaging. Active ROIs are defined as any region with ≥ 50% event frequency increase post-uncaging; see Extended Data Fig. 3j (GABA: *n* = 195 active ROIs; glutamate: *n* = 171 active ROIs from 27 paired FOVs). **k**, Example FOV of ROIs with any baseline events (left) and active ROIs following uncaging (right). Yellow dot = NT uncaging site. **l**, Fraction of ROIs active (*responding*) following both GABA and glutamate uncaging, among all active ROIs for uncaging of either NT (black vertical line; 8.27±1.34%, mean±sem; *n* = 27 paired FOVs). This overlap fraction is compared with a distribution of 10,000 surrogate overlap fractions obtained by choosing an equal number of active ROIs as observed in each FOV, but at random from among all ROIs with any baseline events for GABA and glutamate (grey distribution).

To investigate whether gap junctional coupling mediates these non-cell autonomous Ca^2+^ activity changes after a single point of network stimulation, we either genetically or pharmacologically inhibited gap junctions and measured population-wide network Ca^2+^ responses (Fig. 3c–f). Genetically, we focused on the predominant connexin protein (Cx43) expressed in cortical astrocytes^20,21,45^ (Extended Data Fig. 3a), and decreased Cx43 expression by injecting the astrocyte-specific Cre virus *AAV5-GFAP(0*.*7)-RFP-T2A-iCre*^46^ (and *AAV5-GfaABC1D-GCaMP6f-SV40* to express GCaMP) into Cx43^fl/+^ and Cx43^fl/fl^ mice. Cx43 protein decreases in Cre^+^ cells were confirmed via immunohistochemistry (Extended Data Fig. 3b,c and Supplementary Video 7). After targeting Cre^+^ astrocytes for RuBi-GABA and -glutamate uncaging, population-wide network activity changes were attenuated compared to those observed in wild-type (WT) slices (Fig. 3c, right). While population-wide fluorescence did rise above threshold in some post-stim frames in Cx43^floxed^ and CBX networks with similar onset latencies to WT networks (Fig. 3d and e), the proportion of time population-wide activity remained in an elevated state was significantly reduced in networks with gap junctional inhibition (Fig. 3c and f). Additionally, Cx43^floxed^ networks showed no significant increase in average event frequency, similar to the laser-uncaging controls and receptor-activation controls in slices where GABA_B_R or mGluR were inhibited pharmacologically during uncaging (Extended Data Fig. 3g,h). These results suggest that astrocytic Cx43-based signaling is necessary for the sustained increase in network-level Ca^2+^ activity following NT release elsewhere in the local network. Further, these observations hint that reduced Ca^2+^ signaling in uncoupled astrocyte networks may underlie altered neuronal network activity and deficits in sensory-related behaviors observed in Connexin-deficient mice^47,48^.

We next asked how far NT-induced local network activity extended from the uncaged cell. First, using a Sholl-like analysis (Fig. 3g), we observed event frequency increases as far away as 125– 175μm from uncaging of both GABA and glutamate (Fig. 3h), which extends to the edge of the FOV (Extended Data Fig. 3i). To compare the spatial distribution of these network-level responses between GABA and glutamate, we then analyzed event activity within 20×20μm ROIs in a grid over the FOV (Fig. 3i–k). As in the Sholl-like analysis (Fig. 3h), ROIs with uncaging-driven activity were distributed both near and far from the uncaging site (GABA: 119.9±46.1μm; glutamate: 109.3±49.4μm [mean±SD]) (Fig. 3j). Further, while baseline activity encompasses contiguous, overlapping portions of the astrocyte network (Fig. 3k, left), ROIs exhibiting an event increase after NT uncaging were sparse (Extended Data Fig. 3j) and, critically, exhibit no significant overlap between responses to GABA and glutamate (Fig. 3k, right and l), suggesting that GABA and glutamate do not primarily activate the same regions of the astrocyte network. Together these data show that focal release of NT at a single cortical astrocyte leads to spatially distributed changes in Ca^2+^ activity across an astrocyte network.

### Propagation distinguishes network responses

Because astrocyte events are highly heterogeneous^9^, we next performed an unbiased analysis screen for changes in 16 AQuA-extracted event characteristics from neighboring cells (Extended Data Fig. 4a,b). The most robust and consistent NT-specific changes in neighboring cells were in events exhibiting propagation, with specific directionality toward the pia (Fig. 4a,b, Extended Data Fig. 4b and Supplementary Videos 3–6), which echoed a change we observed above in populations of astrocytes following more widespread NT exposure (Fig. 1h). These propagative events were discrete events contained within individual cells (Fig. 4a) and we did not observe coordinated activity propagating across populations of astrocytes with a visible wavefront (Extended Data Fig. 5a). Since propagative events constituted a small subset of spontaneous *ex vivo* astrocyte Ca^2+^ activity (Extended Data Fig. 5b), we wanted to ensure that they reflected activity that could be observed *in vivo*. To test this, we recorded spontaneous astrocyte Ca^2+^ activity from the same cortical region (V1) in head-fixed mice^5,9^ (Fig. 4c). We focused on spontaneous astrocyte Ca^2+^ activity that occurred when the mouse was stationary, to eliminate stimulus-triggered Ca^2+^ bursts driven by locomotion^9–11,49^. We found a similar fraction of propagative events *ex vivo* and *in vivo* (Fig. 4d), suggesting that this small subset of Ca^2+^ activity may indeed comprise a physiologically relevant population.

**Fig. 4.**
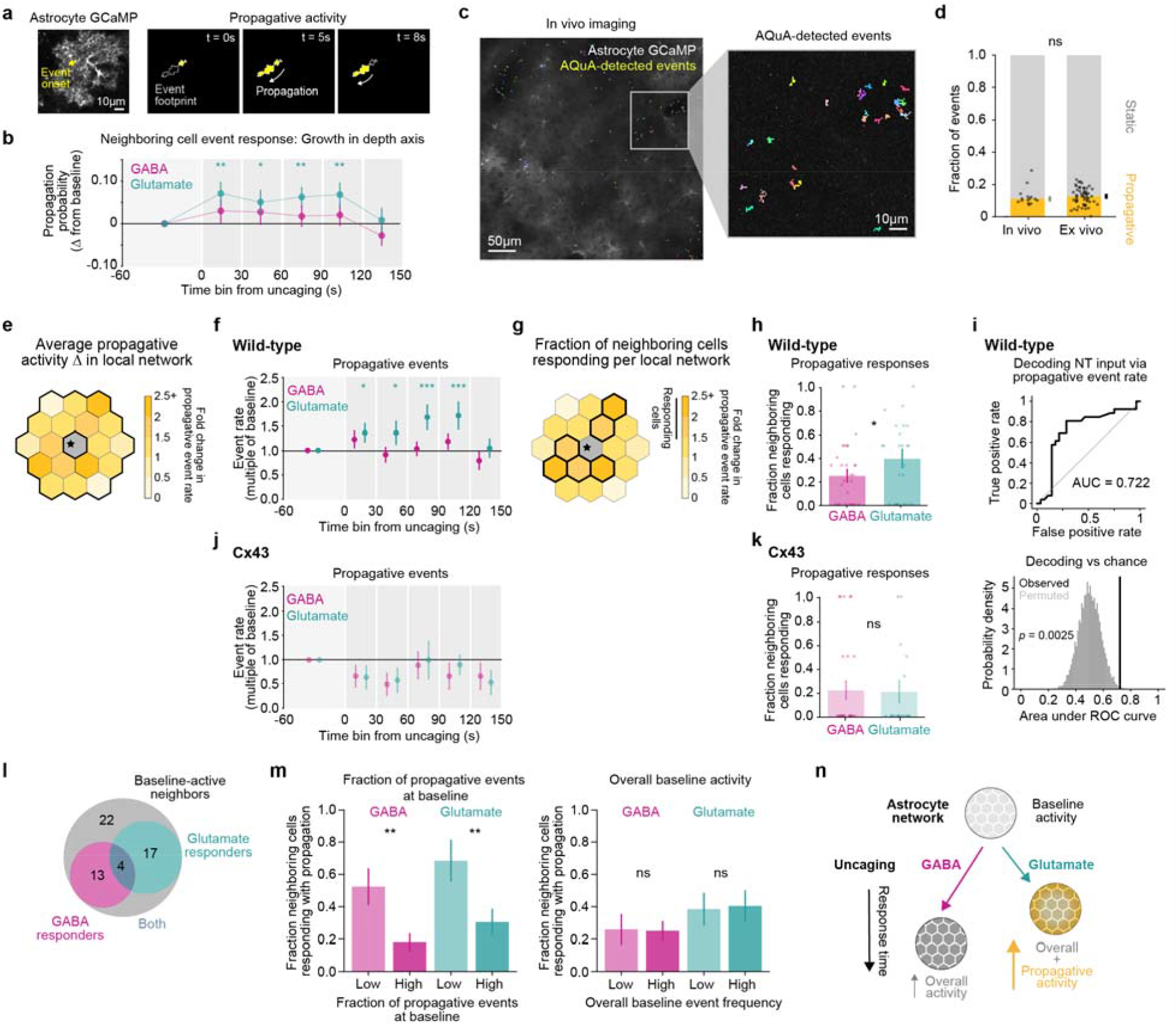
Propagative activity distinguishes astrocyte network responses to GABA and glutamate. **a**, GCaMP6f fluorescence in an individual astrocyte highlighting the initial territory of a propagating event (left). Trajectory of the propagative event over time (3 right panels). The total territory of the event (‘event footprint’) is outlined in grey. The territory of the event at each time point is shown in yellow. **b**, Change in probability of a Ca^2+^ event growing in the depth-axis (toward or away from the pia) among all events from neighboring cells after GABA or glutamate uncaging, relative to 60–0s pre-uncaging. Data shown as overall probability ± standard error determined from hierarchical bootstrapping (Methods; GABA: *n* = 142 cells in 28 FOV; glutamate: *n* = 120 cells in 27 FOV). Two-sided *p*- and *q*-values for changes versus baseline were obtained by circularly shifting each cell’s events in time (Methods; Extended Data Table 8). **c**, *In vivo* 2P image showing expression of astrocyte GCaMP6f in V1. Colored overlay: all AQuA-detected Ca^2+^ events from a 90s stationary period from a 400 x 400μm FOV (left) and 100 x 100μm inset (right). **d**, Fraction of astrocyte Ca^2+^ events that exhibit growth >1μm in any direction (“propagative” events) observed in V1 during stationary wakefulness *in vivo* (left, markers are individual recordings) and at baseline in acute V1 slices (right, markers are individual FOV). Data shown as median across recordings ± standard error via bootstrapping (*in vivo*: *n* = 15 recordings, 5 mice; *ex vivo*: n = 55 recordings, 4 mice). *In vivo* and *ex vivo* setting were compared with a two-sided rank-sum test (*p* = 0.57). **e**, Analysis schematic illustrating average propagative activity change across all neighboring cells in the local network, as in f & j. Note heterogeneity of responses of individual neighboring cells averaged over in f & j. As in Fig. 3, activity from the directly stimulated astrocyte is excluded from all analyses in this figure. **f, j**, Fold-change in rate of propagative Ca^2+^ events among neighboring cells after GABA or glutamate uncaging in acute slices from WT mice (**f**) or Cx43^floxed^ mice (**j**), relative to 60–0s pre-uncaging. Data shown as median across FOVs ± standard error via hierarchical bootstrapping (Methods; *n* for all conditions are in Extended Data Table 9). One-sided *p*- and *q*-values were obtained via circular permutation testing (Methods; Extended Data Table 10). **g**, Analysis schematic illustrating the fraction of neighboring cells per FOV that respond to NT with increases in propagative activity, as in h & k. Note that a subset of neighboring cells in the local network exhibit activity changes following NT uncaging. **h, k**, Fraction of neighboring cells per FOV with ≥ 50% increase propagative Ca^2+^ events (“responding”) after GABA or glutamate uncaging in WT (**h**) or Cx43^floxed^ slices (**k**). Data shown as mean ± sem via hierarchical bootstrapping; dots denote individual FOVs (see Methods; *n* for all conditions are in Extended Data Table 9). Permutation testing was used to compare fraction of cells responding to GABA and glutamate in WT slices (*p* = 0.046) and Cx43^floxed^ slices (*p* = 1.0). **i**, Top: Receiver operating characteristic (ROC) curve of decoding NT identity by thresholding the relative change in propagative event rate from 60–0s pre-uncaging to 0–120s post-uncaging across all neighboring cells in a FOV indicates that propagative events reliably distinguish NT input in WT slices. Bottom: Observed area under the ROC curve = 0.72 ± 0.077 (value ± bootstrapped standard error), compared to a permuted distribution *via* permuting NT labels (*p* = 0.0025, *n* = 55 FOVs). Feature values ≥ threshold were classified as glutamate, and values < threshold as GABA. **l**, Number of neighboring cells responding to one or both NTs with propagative activity increases, among cells with baseline propagative activity (*n* = 56 cells, 24 paired recordings, 7 slices, 4 mice). Permutation testing measures of correlation (Spearman rho, *p* = 0.24) or overlap (Jaccard index, *p* = 0.96) between GABA and glutamate responses does not reject the null hypothesis of independent responses. **m**, Fraction of neighboring cells per FOV with ≥ 50% increase in propagative Ca^2+^ events (“responding”) after GABA or glutamate uncaging in WT slices, split between cells with fraction of propagative events at baseline (left) or overall baseline event rates (right) in the bottom 50% (“Low”, light bars) or top 50% (“High”, dark bars) across all cells (see Extended Data Fig. 5e). Data shown as mean ± sem via hierarchical bootstrapping (*n* in Extended Data Table 11). Response fractions for cells with “Low” and “High” baseline fractions were compared by permuting cells’ baseline propagation fractions for GABA (*p* = 0.0022) and glutamate (*p* = 0.0059); responses for cells with “Low” and “High” overall baseline event rates were compared similarly (GABA: *p* = 1.0; glutamate: *p* = 1.0). **n**, Integrated model of astrocyte network response to GABA and glutamate from data in Fig. 3–4. Astrocyte networks increase general Ca^2+^ activity in response to both NTs, and propagative activity increases specifically in response to glutamate. Network responses to glutamate occur soon after stimulation, while responses to GABA are delayed. For **b & f**: *: *q* < 0.05, **: *q* < 0.01, ***: *q* < 0.001. For **h & m**: *: *p* < 0.05, **: *p* < 0.01.

*Ex vivo*, propagative event frequency specifically increases after glutamate uncaging, in all 30-s time-bins 0–120s post-uncaging across neighboring cells (Fig. 4e, f and Extended Data Fig. 5c), while no changes were observed across neighboring cells after GABA uncaging in these same slices. Indeed, local network responses to glutamate and GABA uncaging can also be distinguished via the fraction of cells exhibiting changes in propagative event frequency (Fig. 4g), in which a higher fraction of astrocytes in each local network respond with increases in propagative activity to glutamate (∼40%) compared to GABA (∼25%) (Fig. 4h and Extended Data Fig. 5d). Further, the NT input received can be accurately decoded via the relative change in propagative event rate/FOV (Fig. 4i). In contrast, a similar fraction of local network astrocytes responds to GABA and glutamate with increased static event frequency (Extended Data Fig. 6). Astrocytes in the local network exhibited significantly higher baseline propagative activity and similar levels of static activity prior to uncaging GABA compared to glutamate (Extended Data Fig. 5g). While this could influence results, these baseline differences likely do not account for the differential network responses to the two NTs because baseline propagative rate is not correlated with the relative post-uncaging propagative event rate (Extended Data Fig. 5h). These results indicate that glutamate and GABA are differentially encoded at the network level by engaging local network astrocytes to differing degrees via Ca^2+^ events that propagate within individual cells (Fig. 4n). Because there are few propagative events at baseline, even a small increase in propagative events following uncaging is a large relative increase in activity, and may thus constitute a salient signal with a high signal-to-noise ratio. This increase in glutamate-driven propagative responses is not observed when uncaging NTs in astrocyte networks with decreased Cx43 expression (Fig. 4j,k), which show significantly lower baseline activity compared to WT networks (Extended Data Fig. 5f). These data suggest that gap junction coupling is a cellular mechanism underlying this NT-specific signal.

Similar to our prior finding that all network-level responses to glutamate and GABA were spatially non-overlapping (Fig. 3k,l), out of astrocytes that responded to either NT, we see few astrocytes responsive via propagative activity increases to *both* NTs (Fig. 4l). In fact, the number of astrocytes responsive to both NTs is not significantly different from chance, indicating that how an astrocyte in the network responds to one NT provides no information about how that same astrocyte will respond to the other NT. Further, when uncaging less glutamate in a different set of local networks (Extended Data Fig. 7a, b), the response profile of an individual astrocyte to three sequential rounds of NT release at the same location was variable (Extended Data Fig. 7c). This was a controlled comparison since average increases in event frequency occurred over a similar time course (Extended Data Fig. 7d) and baseline activity was comparable in astrocytes of the local network across rounds (Extended Data Fig. 7e). Since propagative response to a particular NT does not predict the response to the other NT nor to sequential stimulation by the same NT, we next looked for metrics that instead might predict astrocyte-network responses. Astrocyte Ca^2+^ activity can depend on prior and current Ca^2+^ levels^11,50,51^, which led us to ask whether network-level propagative responses could be predicted based on ongoing network activity. To do so, we asked whether the composition of baseline (1 min) activity in the network could predict the network-level response to uncaging (Extended Data Fig. 5e). Here, we saw that those cells with a higher fraction of propagative events at baseline (relative to all baseline events) were correlated with a lower fraction of astrocytes in each network responding to either GABA or glutamate (Fig. 4m, left, Extended Data Fig. 5i, j). In contrast, overall baseline event rate was not predictive of responses to either NT (Fig. 4m, right). Thus, in addition to differentiating the local astrocyte network response to GABA or glutamate, these data suggest that propagative events can selectively bias the astrocyte network’s subsequent responses to NTs.

## Discussion

Single astrocyte simulation can cause long-lasting changes in neuronal activity and plasticity extending tens to hundreds of microns from the stimulation site^16,17,47^. However, the mechanism(s) that drive distributed effects have been unclear. Here, a brief, spatially restricted NT input leads to long-lasting, network-wide changes in astrocyte Ca^2+^, an effect facilitated by gap junctions. These findings could bridge the spatiotemporal gap between transient, local astrocyte stimulation and sustained, distributed effects on neurons. Still, the full spatial extent of astrocyte network activation remains an open question because astrocyte Ca^2+^ changes extend beyond our FOV.

What might be an effect of restricted NT inputs causing prolonged and distributed responses? For coordinated behavior and learning, neuronal signals are integrated over seconds and minutes^52^. Models of neural integration that solely rely on neuronal activity require fine-tuned positive feedback loops to allow for integration over periods longer than tens of milliseconds^53^. While recurrent neuronal connections enable temporal integration, astrocyte networks provide another possible mechanism to integrate inputs over long time periods^13,54,55^, linking the milliseconds timescale of neurons and the seconds-to-minutes timescales of behavior.

Both GABA and glutamate uncaging led to sustained, far-reaching changes in astrocyte network Ca^2+^ activity, but propagative activity differentiated responses to each NT (Fig. 4n). These propagative events may facilitate integration of information across cellular compartments which could allow for coordinated modulation of groups of nearby synapses^56^ or spatiotemporal integration of inputs across individual cells^51^. Stimulation by glutamate consistently led to greater increases in propagative activity (Fig. 1h and Fig. 4b, f and h), suggesting that cortical astrocytes are more responsive to glutamatergic than GABAergic signaling, as has been described for other brain regions^18^. Heightened astrocyte sensitivity to glutamate may mirror structural organization in the cortex, where astrocyte processes are closer to glutamatergic than GABAergic synapses^57^, potentially reflecting astrocytes’ key role in extracellular glutamate uptake. Since the surface mobility of astrocytic glutamate transporters is dependent on intracellular Ca^2+^ (ref. ^58^), a more robust Ca^2+^ response to glutamate may allow astrocytes to efficiently take up extracellular glutamate by increasing surface mobility of glutamate transporters.

Astrocyte network responses to glutamate and GABA were also context-dependent: responses to both NTs were inhibited specifically by high fractions of propagative events at baseline (Fig. 4m). Thus, as glutamatergic input preferentially recruits propagative events in the surrounding astrocyte network (Fig. 4b,f,h), it could also suppress subsequent responses to NT inputs. This suggests that astrocyte networks implement combinatorial logic, integrating NT inputs across the local network by disseminating information via specific subtypes of Ca^2+^ activity.

While most astrocytes and local networks increase Ca^2+^ in response to NT uncaging, a subset don’t respond to direct or remote uncaging. This heterogeneity may be shaped by the activity state of the astrocyte and connected network during stimulation or by the subcellular location of uncaging. Alternately, only a subset of astrocytes may be equipped to respond to NTs, given the molecular heterogeneity of astrocytes^59,60^. Future experiments imaging astrocyte responses to NTs, followed by spatial transcriptomics, could elucidate how cellular machinery may underlie heterogeneous responses.

Here, astrocytic gap junctions mediate network activity changes, and may also regulate Ca^2+^ activity in individual cells. Molecules, including Ca^2+^ and IP_3_, can diffuse through gap junctions^61,62^. IP_3_ is required for Ca^2+^ release from internal stores^63^ and Ca^2+^ itself regulates Ca^2+^ release from internal stores via calcium-induced calcium release. Here, reduced gap junctional coupling between astrocytes may have altered cytosolic Ca^2+^ and IP_3_ concentrations, which could impact Ca^2+^ release from internal stores and shape Ca^2+^ dynamics within individual cells.

## Methods

### Animals

Experiments were carried out using young adult mice, in accordance with protocols approved by the University of California, San Francisco Institutional Animal Care and Use Committee (IACUC). Animals were housed in a 12:12 light-dark cycle with food and water provided *ad libitum*. Male and female mice were used whenever available. Transgenic mice used in this study were Cx43^fl/fl^ mice^64^ from the Bhattacharya Lab (UCSF, USA) and EAAT2-tdT mice^65^ from the Yang Lab (Tufts University, USA). For *in vivo* imaging, all experiments were performed at the same time each day.

### Surgical procedures

For viral expression for *ex vivo* experiments, neonatal Swiss Webster or C57Bl/6 (P0–3) mice were anesthetized on ice for 3 min before injecting viral vectors (*AAV5*.*GfaABC*_*1*_*D*.*GCaMP6f*.*SV40* [Addgene, 52925-AAV5], *AAV9*.*hGfap*.*pinkFlamindo, pENN*.*AAV9*.*Gfap*.*iGluSnFr*.*WPRE*.*SV40* [Addgene, 98930-AAV9], or *AAV5*.*GFAP(0*.*7)*.*RFP*.*T2A*.*iCre* [Vector Biolabs, 1133]). Pups were placed on a digital stereotax and coordinates were zeroed at lambda. Four injection sites in a 2 × 2 grid pattern over V1 were chosen. Injection sites were 0.8–0.9 mm and 1.6–1.8 mm lateral, and 0 and 0.8–0.9 mm rostral. At each injection site, 30–120 nl of virus were injected at a rate of 3 nl/s at two depths (0.1 mm and 0.2 mm ventral/below pia) using a microsyringe pump (UMP-3, World Precision Instruments).

For viral expression for the *in vivo* experiments, adult C57BL/6 mice were administered dexamethasone (5mg/kg, s.c.) >1 hour before surgery, and anesthetized using 1.5% isoflurane (Patterson Veterinary Supply, 78908115). After hair removal and three alternating swabs of 70% ethanol (Thermo Fisher Scientific, 04-355-720) and Betadine (Thermo Fisher Scientific, NC9850318), a custom-made titanium headplate was attached to the skull using cyanoacrylate glue and C&B Metabond (Parkell, S380). A 3mm craniotomy was made over the right visual cortex. Virus was injected at two sites in right visual cortex at coordinates centered on +2.4mm and +2.7mm medial/lateral, +0.35mm and +0.65mm anterior/posterior and -0.3mm dorsal/ventral from lambda. 300nL of *AAV5*.*GfaABC*_*1*_*D*.*GCaMP6f*.*SV40* (Addgene, 52925-AAV5) was injected at each site through a glass pipette and microsyringe pump (UMP-3, World Precision Instruments). After allowing at least ten minutes for viral diffusion, the pipette was slowly withdrawn and a glass cranial window implanted using a standard protocol.

### *Ex vivo* two-photon (2P) imaging and uncaging

Coronal, acute V1 slices (400-μm thick) from P28–32 (bath-application) and P27–42 (uncaging) mice were cut with a vibratome (VT 1200, Leica) in ice-cold slicing solution containing (in mM) 27 NaHCO_3_, 1.5 NaH_2_PO_4_, 222 sucrose, 2.6 KCl, 2 MgSO_4_, 2 CaCl_2_. Slices were transferred to pre-heated, continuously aerated (95% O_2_/5% CO_2_) standard artificial cerebrospinal fluid (ACSF) containing (in mM) 123 NaCl, 26 NaHCO_3_, 1 NaH_2_PO_4_, 10 dextrose, 3 KCl, 2 MgSO_4_, 2 CaCl_2_. Younger mice were sliced in the same solutions for GCaMP bath application of LY379268 and Baclofen (P20–25), Pink Flamindo (P20–22), and GluSnFR (P14–17). Slices were kept at room temperature until imaging. Bath-application experiments were performed at room temperature and 2P uncaging experiments were performed at 29°C using an in-line heater (TC-324B and SH-27B, Warner Instruments). To block neuronal action potentials during all slice imaging experiments, TTX (1 μM) was added to the ACSF > 10 min before imaging and remained in the circulating bath for the duration of the experiments.

Images were acquired on an upright microscope (Bruker Ultima IV) equipped with two Ti:Sa lasers (MaiTai, SpectraPhysics). Laser beam intensities were modulated using two independent Pockels cells (Conoptics) and images were acquired by scanning with linear galvanometers. Images were acquired with a 16×, 0.8 N.A. (Nikon) or a 40×, 0.8 N.A. (Nikon) water-immersion objective via photomultiplier tubes (Hamamatsu) using PrairieView (Bruker) software. For GCaMP imaging, 980 nm excitation and a 515/30 emission filter were used. For RFP imaging, 980 nm excitation and a 605/15 emission filter were used. For Pink Flamindo and Alexa Fluor 594 imaging, 1040 nm excitation and a 605/15 emission filter were used. Images were acquired at 1.42 Hz frame rate, 512 × 512 pixels and 0.64–1.61 μm/px resolution. For GluSnFR imaging only, images were acquired at 6.21 Hz frame rate, 200 × 200 pixels and 0.64 μm/px resolution, with 980 nm excitation and a 515/30 emission filter.

For bath-application experiments, a 5-min baseline was recorded to monitor spontaneous activity, after which receptor agonists were added along with a fluorescent dye (Alexa Fluor 594 Hydrazide) to assess the time at which drugs reached the imaging field/recording chamber (except for Pink Flamindo due to spectral overlap). An ACSF washout period (> 10 min), followed by a TTX incubation period (>10min), occurred between trials when imaging the same slice sequentially for bath-application of different receptor agonists or uncaging of different RuBi-subtypes. To account for any changes resulting from prior agonist exposure or uncaging, we alternated the order of agonists between concentrations or RuBi-subtypes between slices.

For simultaneous 2P imaging and uncaging, a second Ti-Sa laser beam was tuned to 800 nm and controlled using an independent set of linear galvanometers from those used for scanning. Laser beam intensity was modulated using an independent Pockels cell (Conoptics) to achieve a power measurement of ∼2–8 mW at the slice. The beam paths for imaging and uncaging were combined after the linear galvanometers using an 855-longpass dichroic mirror (T855lpxr, Chroma). The uncaging laser was calibrated each experimental day by burning spots into a fluorescent slide. RuBi-compounds (300 μM) and TTX (1 μM) were added to the ACSF >10 minutes before imaging each slice. Fields of view (FOV) were chosen based on the location of GCaMP expression, which was often biased to/brighter in deeper cortical layers (distance of FOV from pia: 615 ± 196μm [mean ± SD, n = 121 FOV]). Prior to imaging at each FOV, a 60-s period was recorded to identify potential uncaging sites. Areas of GCaMP expression that exhibited moderate levels of spontaneous Ca^2+^ activity were chosen as uncaging sites. For FOVs with sequential GABA/glutamate uncaging, a continuous 5-min recording was used to monitor activity in each FOV. For FOVs with three sequential rounds of glutamate uncaging, a continuous 12.5-min recording was used to monitor activity in each FOV. Each recording began with a 2.5-min baseline period and at the 2.5-min mark, neurotransmitter was uncaged with 10 x 100 ms pulses, 100 ms apart. Sequential recordings of GABA/glutamate uncaging within the same FOV were separated by > 20min. Rounds of sequential glutamate uncaging were separated by ≥ 25min. Voltage from the uncaging laser Pockels cell was recorded to mark the time of uncaging pulses. Because RuBi-GABA and RuBi-glutamate are light-sensitive, care was taken to ensure experiments were carried out in minimal light. The computer screen and red-shifted headlamp were covered with two layers of red filter paper (Roscolux #27 filter, Rosco) and all indicator lights on equipment were covered.

### *In vivo* 2P imaging

*In vivo* 2P imaging was performed on the same microscope as *ex vivo* imaging, via a Nikon 16x, 0.8 N.A. water-dipping objective with a 2x-optical zoom (frame rate: 1.7Hz, FOV: 412μm^2^, resolution: 512×512 pixels). Animals were given > 1 week after surgery for recovery and viral expression. They were then habituated on a custom-made circular running wheel over at least two days, and for a cumulative time of at least 2.5 hours, before recording. After habituation, mice were head-fixed on the wheel and movements were recorded by monitoring deflections of colored tabs on the edge of the wheel using an optoswitch (Newark, HOA1877-003). To compute wheel speed, a detected break in the optoswitch circuit was determined when the absolute value of the derivative of the raw voltage trace was at least 2 standard deviations above the mean. For recordings with little movement (std < 0.1), this threshold generated false positives, so a set threshold of 0.1 was used. The number of breaks in the optoswitch circuit per second was then calculated, and using the circumference and number of evenly spaced colored tabs at the edge of the wheel, the wheel speed was determined and used for all subsequent analyses using speed. Movement periods were defined by wheel speed ≥ 10 cm/s and movement bouts that were separated by ≤ 2 s were considered one event. To ensure that movement related dynamics were not included in stationary analysis, data was excluded from < 10 s around identified movement periods. GCaMP was imaged with 950nm excitation light and a 515/30 emission filter. Recordings lasted 30 minutes.

### *Ex vivo* pharmacology

The following concentrations of each pharmacological reagent were used for experiments as indicated in the text: Tetrodotoxin Citrate (TTX, 1 μM, Hello Bio); Carbenoxelone disodium (CBX, 50 μM, Tocris Bioscience); R(+)-Baclofen hydrochloride (5–100 μM, Sigma-Aldrich); (1S,3R)-ACPD (t-ACPD, 5–100 μM, Tocris); LY 379268 disodium salt (100 μM, Tocris); Alexa Fluor 594 Hydrazide (0.1–2 μM, ThermoFisher Scientific); RuBi GABA trimethylphosphine (RuBi-GABA-Pme_3_, 300 μM, Tocris); RuBi-Glutamate (300 μM, Tocris); CGP 55845 hydrocholoride (10 μM, Tocris); and LY 341495 (10 μM, Tocris).

### Immunohistochemistry and image quantification

After recording, slices from 2P imaging experiments were immersed in 4% PFA for 30 min and switched to 30% sucrose for one day at 4°C before being embedded in OCT and stored at -80°C. Slices were re-sectioned coronally at 40 μm on a cryostat and then stored in cryoprotectant at - 20°C until staining. For immunohistochemistry, sections were washed three times in 1X PBS for 5 min and permeabilized for 30 min with 0.01% Triton-X in 1X PBS. Sections were next blocked with 10% NGS (Abcam) for 1 h and incubated overnight with primary antibodies at 4°C in 2% NGS. The next day, they were washed three times in 1X PBS before incubating with secondary antibodies for 2 h at room temperature. Sections were washed three times in 1X PBS for 5 min before being mounted on slides with Fluoromount-G (SouthernBiotech).

To validate reduction of Connexin 43 (Cx43) protein in astrocytes transduced with AAVs to express GCaMP-GFP and Cre-RFP, primary antibodies for α-connexin-43 (1:1500, rabbit, Sigma-Aldrich), α-GFP (1:3000, chicken, Abcam), and α-mCherry (1:2000, rat, Thermo Fisher Scientific) in 2% NGS were used. Secondary antibodies include α-rabbit Alexa Fluor 405, α-chicken Alexa Fluor 488, and α-rat Alexa Fluor 555 (all Thermo Fisher Scientific), which were all used at 1:1000 dilution. 60x multi-channel z-stack images were acquired on a CSU-W1 Spinning Disk Confocal (Nikon) from V1 in which AAVs were injected. To quantify loss of Cx43 in RFP^+^ and RFP^-^ astrocytes, Fiji (ImageJ) was used. Through batch processing, cell maps were created through a semi-automated pipeline to segment astrocytes, with *post hoc* ROI adjustments for vasculature artifacts. Multi-channel z-stacks were split into 405, 488, and 555 channels, and unstacked into sequential 8-bit z-plane images. For each z-plane, RFP^+^ and RFP^-^ astrocytes were detected using a Gaussian blur (sigma = 3), thresholding using the Phansalkar method (radius = 1000), and applying ImageJ’s “Analyze Particles” (size > 175 μm^2^, circularity = 0–0.60) to outline ROIs using the wand tool. Corresponding Cx43 images were binarized and the Fiji plugin SynQuant^66^ was used to detect Cx43 puncta number within each RFP^+^ and RFP^-^ astrocyte in a z-plane’s cell map. Puncta counts were normalized to astrocyte area, and the normalized count from each z-stack was averaged for each slice.

### 2P image and data analysis

Individual-astrocyte cell maps for time series images were created in Fiji using the following process: For each FOV, an 8-bit z-projection of the time series was created. The z-projection was binarized using the ‘Auto Local Threshold’ feature, using the Niblack method and a radius of 30 or 75, for 16× and 40× images, respectively. Cell maps were drawn on binarized images using a combination of the Lasso and Blow Tool and freehand drawing tool in Fiji, and verified on the z-projected image. Cell maps were also verified against a static indicator of astrocyte morphology when available (EAAT2-tdT^+^ mice for bath-application of LY379268 and Baclofen; GFAP(0.7)-RFP-T2A-iCre in Cx43^floxed^ mice). To load cell masks into AQuA, regions were saved to the ROI manager and filled in with a color. The regions were projected onto a black image the same size as the original (512 × 512 pixels). The overlay of regions was flattened, converted to an 8-bit image, and saved as a tiff. For the 12.5-min recordings with sequential rounds uncaging glutamate, drift of the slice in X and Y was corrected post-hoc using moco^67^.

#### AQuA

GCaMP and GluSnFR 2P image sequences were analyzed using AQuA^9^ and custom MATLAB code (MATLAB R2018b). Signal detection thresholds were adjusted for each video to account for differences in noise levels after manually checking for accurate AQuA detection. Cell maps were loaded into AQuA to define cells consistently over multiple time-series featuring the same FOV. For all bath-application experiments, the direction of pia was marked as anterior. For 2P uncaging experiments, the uncaging site was marked as a 3 × 3-pixel landmark.

#### Bath-application event-based analysis

For Baclofen and t-ACPD Ca^2+^ imaging experiments, *Event count per frame* was quantified by counting all AQuA-detected events, new or ongoing, in each frame (Fig. 1c). *Percent of field active* values were calculated by recording the number of active pixels in each frame, as determined by the frame-by-frame footprints of AQuA-detected events. These values were normalized by total number of active pixels in the recording and multiplied by 100. For the *Percent of field active* dose-response curve (Fig. 1e), the percent of field active values from all frames within the chosen timepoints were averaged by slice. *Event propagation* was calculated by summing the growing propagation from all cardinal directions, using the AQuA feature *propGrowOverall*. For dose-response curves for discrete event features (area, duration and propagation [Fig. 1f–h]), all detected Ca^2+^ events within the chosen timepoints were averaged by slice.

The frame the agonist entered the recording chamber was estimated using fluorescence from Alexa Fluor 594 (0.1–2 μM) added to the ACSF reservoir along with agonist. The frame agonist entered the recording chamber was estimated using the maximal curvature method on frames 1– 600 of the raw Alexa Fluor 594 fluorescence trace. The *maximum curvature method*^68^ defines the onset fluorescence changes as the point of maximum curvature during the rising phase of the signal. To identify this point, traces were fit using a modified Boltzmann’s sigmoidal equation:

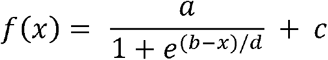

where *a* is the difference between the minimum and the maximum fluorescence, *b* is the inflection point, *c* is the baseline fluorescence and *d* is the slope, using a nonlinear least squares algorithm (Levenberg-Marquardt) in MATLAB (Mathworks). Next, the frames of maximum curvature were calculated by setting the fourth derivative of the fitted curve equal to zero and solving for its three solutions. The earliest frame identified out of these three solutions was recorded as the onset frame.

#### Bath-application ROI-based analysis

Pink Flamindo and GCaMP imaging experiments were analyzed using ROI-based approaches in Fiji. To identify responding cells in Pink Flamindo experiments (Extended Data Fig. 1j), sigmoidal curves were fit to ΔF/F traces using the modified Boltzmann’s sigmoidal equation detailed above. Cells were defined as “responding” if the difference between the minimum and maximum values of the fit curve (*a* in the Boltzmann’s sigmoidal equation) > baseline noise (3 SD of baseline fluorescence). Responding cells were defined as “increasing” if *f*(*x*_start_) < *f*(*x*_end_) and decreasing if *f*(*x*_start_) > *f*(*x*_end_).

To identify fluctuations in Pink Flamindo and GCaMP fluorescence (Extended Data Fig. 1k), peaks were detected from ΔF/F traces from individual cells. Peaks were counted if they were 3 SD above the mean baseline fluorescence, had a minimum peak width of 5 frames and a minimum distance of 10 frames between detected peaks. The baseline period was defined as all frames before the frame of agonist entry. For GCaMP, all astrocytes exhibiting ≥ 1 AQuA-detected event during the 10-min recording were run through peak finding. For Pink Flamindo, all detected astrocytes were run through peak finding.

For GCaMP experiments, the frame agonist entered the recording chamber was estimated using the fluorescence from Alexa Fluor 594 (0.1–2 μM) added to the ACSF reservoir along with agonist. Time of agonist entry in the recording chamber was estimated by identifying the first frame Alexa Fluor 594 fluorescence reached ≥ 3 SD above baseline mean (frames 1–300); only frames > 375 were considered for evaluation of exceeding the threshold. For Pink Flamindo experiments, dye was not added with agonist to avoid spectral overlap. Time of agonist entry in the recording chamber was estimated by adding 90 frames (the average number of frames for ACSF to travel from the reservoir to the recording chamber) to the frame agonist was added to the reservoir of ACSF

#### 2P uncaging event-based analysis

Individual astrocytes were excluded from analyses (Fig. 2–4, Extended Data Fig. 2–7) if the baseline event rate changed significantly. Changes in baseline event rate for each cell were determined by performing Poisson regression of events in 1-s bins during the period from 90–10s pre-uncaging. Cells with a regression coefficient with p < 0.1 at baseline and with > 5 AQuA-detected events throughout the recording were excluded from all analyses, except for Extended Data Fig. 7d RuBi-glutamate uncaging control. ∆F/F values in raster plots (Fig. 2h and 3c) were calculated using the AQuA output *dffMatFilter(:*,:,*2)*, the ∆F/F traces from events after removing the contributions from other events in the same location. For the Sholl-like analysis (Fig. 3h), events were sorted into 50μm bands radiating out from the uncaging site based on the minimum distance between an event and the uncaging site at event onset (using the AQuA output *ftsFilter*.*region*.*landmarkDist*.*distPerFrame*). In order to categorize events as propagative or static (Fig. 4d–m and Extended Data Fig. 5b–j, 6 and 7c), the total propagation distance of each event was computed by summing the growing propagation from all cardinal directions, using the AQuA feature *propGrowOverall*. Events were categorized as propagative if the total propagation distance > 1μm.

### Statistics for Fig. 1–3 and associated Extended Data Figures

All statistical tests used and exact n values can be found for each figure in the corresponding figure legend. Adjustments for multiple comparisons using Bonferroni-Holm correction were implemented using fwer_holmbonf^69^. Significance levels defined as the following: ns: *p* ≥ 0.05, *: *p* < 0.05, **: *p* < 0.01, ***: p < 0.001.

#### Permutation testing

Statistical significance for time-series (t-series) data was computed using permutation testing with custom-written code in MATLAB. 10,000 permutations were run and one- or two-sided *p*-values for each time point were calculated. *p*-values were corrected for multiple comparisons using the Benjamini-Yekutieli procedure (implemented using ref. ^70^) with a False Discovery Rate (FDR) ≤ 0.05.

Data were shuffled/permuted in the following way: To test change in event number/cell (Fig. 1c, Extended Data Fig. 2b and 3g,h), events were shuffled independently for each active cell (≥ 1 AQuA-detected event) in each t-series. For each active cell, events were randomly placed in time bins spanning the duration of the recording (time bins = 60s [Fig. 1c] and 30s [Extended Data Fig. 2b and 3g,h]) and the change in number of events/time bin was calculated as for the experimental data. Permuted changes in event number/cell were averaged across active cells in each t-series and across all t-series to obtain the permuted mean for one round of permutation testing.

To test change in event number/ band (Fig. 3h), permutation tests were run separately for each band and events were shuffled independently for each t-series. For each t-series, events from the tested band were randomly placed in 30 s time bins spanning the duration of the recording, and the change in event number/30 s was calculated as for the experimental data. Permuted changes in event number/30 s were averaged across all t-series to obtain the permuted mean for one round of permutation testing. To test magnitude of change in experimental data versus permuted data, two-sided *p*-values were calculated as:

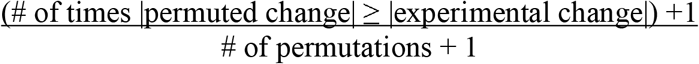

For testing increases in ∆F/F (Extended Data Fig. 1d), frames were shuffled independently for each t-series. For each t-series, the average ∆F/F/frame from active regions (≥ 1 AQuA-detected event in either condition [baclofen or t-ACPD]) was calculated, the frame order was shuffled, and the mean ∆F/F/30s was calculated. Permuted mean ∆F/F was averaged across all t-series to obtain the permuted mean for one round of permutation testing. To test magnitude of increases in experimental data versus permuted data, one-sided *p*-values were calculated as:

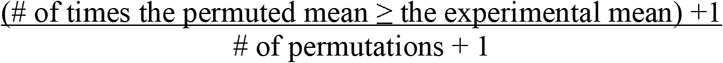

### Statistics for Fig. 3i–l, Fig. 4, and associated Extended Data Figures

#### 2P uncaging grid-based ROI analysis

Grid-based regions of interest (ROIs) were determined by dividing the 300 x 300 μm imaging field into a uniform 20 x 20 μm grid (Fig. 3i–l). Each identified Ca^2+^ event was assigned to the ROI in which the centroid of its spatial footprint was located. ROIs with any baseline events were identified as ROIs with ≥1 events in the baseline window 60–0s before uncaging. “Active” ROIs for each NT were identified as ROIs with a ≥ 50% increase in event rate in the window 0–120s after uncaging for that NT, as compared with the rate during the baseline window. Active ROIs were a subset of ROIs with baseline events, as the relative increase in event rate is not defined when there are no baseline events, which results in division by 0. The distance from the uncaging site to each active ROI was determined using the Euclidean distance between the uncaging site, at (0, 0), and the center of each grid ROI (Fig. 3j).

The fraction of overlap (*i*.*e*., Jaccard index) *O*_*i*_ between active ROIs for GABA and glutamate were determined for the *i*th field of view by

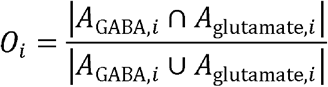

where *A*_GABA,*i*_ and *A*_glutamate,*i*_ are the sets of active ROIs for GABA and glutamate, respectively. The overall fraction of overlap *O* between active ROIs for GABA and glutamate was computed as the mean of the individual *O*_*i*_ (Fig. 3l).

To determine if the observed fraction of overlap was expected due to chance, a distribution of *N* = 10,000 surrogate fractions of overlap was computed. The *k*th surrogate value, *Õ*^(*k*)^ was computed as above, but replacing, for each NT, the set of active ROIs *A*_NT,*i*_ with a new set, 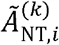, which was chosen as a random subset of size |*A*_NT,*i*_| of the set of ROIs with any baseline events for that NT. The *p*-value for this comparison was estimated^71^ as

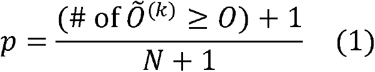

#### Propagation probability (Fig. 4b)

Each Ca^2+^ event was identified as “growing in the depth axis” if the frontier of that event’s spatial footprint extended over time either toward the pia or away from the pia, as determined by the posterior and anterior component of the *propGrowOverall* metric computed *via* segmentation by AQuA^9^.

The probability of events growing in the depth axis was computed separately for recordings of GABA and glutamate uncaging within each examined time window. Probabilities were estimated for the baseline window of 60–0s before uncaging, as well as in nonoverlapping 30s bins ranging from 0–150s post-uncaging, by computing the fraction of events that were identified as growing in the depth axis among all events from all recordings within the relevant time window. The change in the probability of events growing in the depth axis was then estimated for each bin as the difference between the fraction of events growing in the depth axis for that bin versus for the baseline period.

To empirically determine the distribution of each of these estimators, we performed this same procedure for estimating the probability of events growing in the depth axis for each NT and time bin on surrogate data generated by hierarchically bootstrapping Ca^2+^ event data, where the hierarchy was sampled cells within sampled recordings (i.e., all events for an individual cell-recording always remained together); this procedure was repeated 10,000 times for each bin. Standard errors were computed as the standard deviation of these empirical distributions.

To determine the probability of observing effects this large under a null hypothesis of no effect of time on the probability of events growing in the depth axis, we computed the distribution of the estimator under an imposed condition in which the overall temporal structure of astrocyte Ca^2+^ events was disrupted. To do this, we performed the same procedure as above for estimating the probability of events growing in the depth axis for each bin, but on surrogate data generated by circularly shifting the timing of each individual cell’s Ca^2+^ events from 90s before to 150s after uncaging by its own independent, uniform random shift between 0s and 240s; this procedure was repeated *N* = 10,000 times for each bin. As it was unknown whether event propagation would increase or decrease post-uncaging, two-sided *p*-values were estimated^71^ as

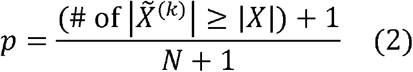

where *X* denotes the actual observed value of the estimator, and each 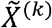 is the value of the estimator computed from the *k*th shifted dataset. These *p*-values were adjusted across tested time bins and NTs using the Benjamini-Hochberg procedure to obtain *q*-values, as implemented in statsmodels 0.12.2 (ref ^72^).

#### Event feature changes (Extended Data Fig. 4a,b)

Each Ca^2+^ event is assigned several metrics by AQuA-segmentation^9^, including size (*area, perimeter, circMetric* [circularity, based on area and perimeter]), amplitude (*dffMax*), and dynamics (*rise19* [rise time], *fall91* [fall time], *decayTau* [decay time constant], *width11* [duration]). For each non-propagation metric, the mean metric value among events was computed separately for recordings of GABA and glutamate uncaging for the baseline window 60–0s before uncaging, as well as in nonoverlapping 30s bins from 0–150s post-uncaging. For each bin, the ratio of that bin’s mean metric value to the baseline mean metric value was computed.

AQuA metrics also capture information about events’ directional propagation. Each Ca^2+^ event was identified as “growing” or “shrinking” in each cardinal direction if the frontier of that event’s spatial footprint extended or receded, respectively, over time in that direction, as determined by the components of the *propGrowOverall* and *propShrinkOverall* metrics. For each propagation metric, the change in the probability of events growing or shrinking in each axis was computed separately for recordings of GABA and glutamate uncaging within each examined time window, as above in *Propagation probability*, but using the “growing” or “shrinking” identifiers for each cardinal direction.

To empirically determine the distribution of each of these estimators (*i*.*e*., binned post/baseline ratio for non-propagation metrics, binned change in growing/shrinking probability for propagation metrics), we performed the same procedures for computing each metric’s relevant estimators for each NT and time bin outlined above on 10,000 surrogate datasets generated by hierarchically bootstrapping Ca^2+^ event data, as described in *Propagation probability*. Standard errors were computed as the standard deviation of these empirical distributions.

To determine the probability of observing effects this large under a null hypothesis of no effect of time on the probability of events growing in the depth axis, we computed the distribution of each estimator under 10,000 realizations of an imposed condition in which the overall temporal structure of astrocyte Ca^2+^ events was disrupted by randomly circularly shifting each cell’s Ca^2+^ events, as described in *Propagation probability*. As it was unknown whether event propagation would increase or decrease post-uncaging, two-sided *p*-values were estimated using equation (2) above^71^. These *p*-values were adjusted across tested time bins and NTs using the Benjamini-Hochberg procedure to obtain *q*-values, as implemented in statsmodels 0.12.2 (ref. ^72^).

#### Comparison of in vivo and ex vivo event propagation (Fig. 4d)

Events were categorized as propagative or static, as outlined above in the *2P uncaging event-based analysis* section. The fraction of propagative events observed *in vivo* and *ex vivo* was calculated using baseline events. Ca^2+^ events in *in vivo* recordings were labeled as “baseline events” if they occurred during periods when the mouse was stationary, as outlined above in the *in vivo 2P imaging* section. Ca^2+^ events in *ex vivo* recording were labeled as “baseline events” if they occurred in neighboring astrocytes (*i*.*e*. cells not directly stimulated by NT) during the 60–0s before NT uncaging.

To determine the distribution of the two median propagative event fractions empirically, we computed the medians of 10,000 bootstrapped samples of the per-recording fractions for each setting. Standard errors for each statistic were determined from the standard deviations of these empirical distributions.

#### Computing rate changes for propagative and static events (Fig. 4f, j and Extended Data Fig. 6b–c)

The overall rates of propagative and static events for neighboring astrocytes were computed separately for recordings of GABA and glutamate uncaging.

For each event class (*i*.*e*., propagative and static events), for each recording, the event rate was computed in each time window as the total number of events from all neighboring cells in that recording in the given time window divided by the duration of that time window. These recording-level rates were computed for the baseline window of 60–0s before uncaging and in nonoverlapping 30s bins ranging from 0–150s post-uncaging. For each recording, the relative rate of propagative and static events was computed for each time bin as the ratio of the event rate for the given event class in that time bin divided by the corresponding event rate in the baseline window. For each time bin, the overall relative rate was estimated as the median of the per-recording relative rates in that time bin.

To determine the distribution of each of these relative rate estimators empirically, we performed this same procedure for estimating relative event rates on surrogate data generated by hierarchically bootstrapping Ca^2+^ event data 10,000 times for each bin (as above in *Propagation probability*). Standard errors were computed as the standard deviation of these empirical distributions.

To determine the probability of observing effects this large under a null hypothesis of no effect of time post-uncaging on the rate of astrocyte Ca^2+^ events, we computed the distribution of the relative rate estimators under an imposed condition in which the overall temporal structure of astrocyte Ca^2+^ events was disrupted *via* a random circular shift of the events in each cell, as above in Fig. 4b; this procedure was repeated *N* = 10,000 times for each bin. Motivated by results in bath application experiments above demonstrating robust aggregate astrocyte Ca^2+^ activity increases in response to agonism of glutamate receptors (Fig. 1h), one-sided *p*-values were estimated from these permuted datasets, as in equation (1) above. These *p*-values were adjusted across tested time bins and NTs using the Benjamini-Hochberg procedure to obtain *q*-values, as implemented in statsmodels 0.12.2 (ref. ^72^).

#### Determining responding cells based on static and propagative events (Fig. 4h,k and Extended Data Fig. 6e–f)

The overall rates of propagative and static events were computed for each neighboring astrocyte, with paired measurements made for recordings of GABA and glutamate uncaging. For each neighboring astrocyte, for each event class (*i*.*e*. propagative and static events), the event rate was computed in each time window as the total number of events from that cell in the given time window divided by window’s duration (baseline window: 60–0s before uncaging, response window: 0–120s after NT-uncaging; Extended Data Fig. 5c). Relative event rates were calculated as for Fig. 4f, j and Extended Data Fig. 6b–c above. Cell-recording combinations with zero events of a given type in the baseline window were excluded for computation of relative rates of propagative (GABA: 36 cell-recordings [26.7% of total]; glutamate: 37 [32.2%]) and static (GABA: 0; glutamate: 0) events, as the relative rate would require a division by zero and be undefined in those cases. Astrocytes were identified as “responders” with a particular event type (*i*.*e*., static or propagative) to GABA or glutamate if their relative rate of that type of event was ≥ 1.5 for the corresponding NT uncaging recording (Extended Data Fig. 5d). The fraction of astrocytes that were responders was computed for each individual recording, as well as the overall fraction of responders across all recordings for each NT.

To determine the distribution of these overall responder fractions, we performed this same procedure for estimating relative event rates on surrogate data generated by hierarchically bootstrapping Ca^2+^ event data 10,000 times (as above in *Propagation probability*). Standard errors were computed as the standard deviation of these empirical distributions.

To determine whether there were significant differences between the overall responder fractions for GABA and glutamate, we computed the distribution of the difference between these two fractions under an imposed condition in which there was no systematic difference between GABA and glutamate. To do this, we performed the same procedure as above for estimating the difference between the overall responder fractions for “GABA” and “glutamate”, but on surrogate data generated by, for each cell, swapping the labels for “GABA” and “glutamate” responses from that in the experimental data with probability 1/2; this procedure was repeated 10,000 times. As it was unknown *a priori* whether GABA or glutamate would have a higher fraction of responder cells, a two-sided *p*-value was estimated as in equation (2) above.

#### Decoding NT identity from propagative event responses (Fig. 4i)

To quantify the extent to which the observed difference in propagative event responses to uncaged glutamate and GABA enabled reliable identification of NT identity on a trial-by-trial basis, we built a simple classifier that took as input a single value, the relative change in propagative event rate across a FOV in the window 0–120s post-uncaging relative to the window 60–0s pre-uncaging, and classified that FOV as responding to glutamate if the value was ≥ a set threshold, and GABA if the value was < the threshold. To evaluate this classifier’s performance, we built a receiver operating characteristic (ROC) curve by varying the classification threshold across the entire domain of the feature, and at each value of the threshold, computing the empirical true positive rate and false negative rate of the classifier. With the threshold fixed in the ROC analysis, the classifier did not have any remaining free parameters, so did not need to be trained on data and was therefore not a function of any of the data, obviating the need for cross-validation. We computed the area under the ROC curve (AUC) using the trapezoidal rule. To determine the distribution of the observed AUC statistic, we performed this same analysis on 10,000 surrogate datasets generated by bootstrapping (*i*.*e*., resampling FOVs with replacement). To determine whether the observed AUC statistic was above 0.5 (indicating completely non-informative decoding) to a degree greater than expected by chance alone, we performed this same analysis on 10,000 surrogate datasets generated by permuting the NT labels.

#### Determining correlations between GABA and glutamate responses (Fig. 4l)

To determine whether individual cells’ responses to GABA and glutamate—as determined in 4h above—were correlated, we computed the Spearman p between the binary paired responses to GABA and glutamate across cells which could be assessed in both conditions (*i*.*e*., had > 0 propagating baseline Ca^2+^ events in both recordings) using SciPy 1.6.2 ^73^. To determine the probability of observing a correlation at least this large under a null hypothesis of independence between cells’ responses for GABA and glutamate, we computed Spearman p on surrogate data in which the identities of the cells’ responses to GABA and glutamate were independently permuted; this procedure was repeated 10,000 times. To maintain the ability to identify correlation or anticorrelation, we estimated a two-sided *p*-value from these surrogate values, as in equation (2).

To complement this analysis, we computed the fraction of overlap (*i*.*e*., Jaccard index) between the sets *C*_GABA_ and *C*_glu_ of cells that were responders to GABA and glutamate, respectively:

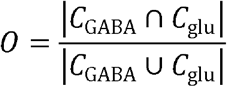

This statistic is larger when the fraction of overlap between responders for the two neurotransmitters is larger. To determine the probability of observing an overlap at least this large under a null hypothesis of independent responses for GABA and glutamate, we computed this same statistic, but on 10,000 permuted surrogate datasets, as above. To determine significant overlap, we estimated a one-sided *p*-value from these surrogate values, as in equation (1).

##### Segregating responding cells based on baseline propagation (Fig. 4m)

For each neighboring astrocyte with propagative events during the baseline period of 60–0s pre-uncaging, we computed the fraction of baseline events that were propagative (# propagative baseline events / # all baseline events). Separately for GABA and glutamate, we used the propagative fraction across all given astrocytes to define the threshold fraction of baseline propagative activity, *f*_50_, as the 50^th^ percentile of all observed values; cells with fractions < *f*_50_ were said to have “low baseline propagation”, while cells with fractions ≥ *f*_50_ were said to have “high baseline propagation” (Extended Data Fig. 5e, top). The fraction of astrocytes that were responders with propagative events to GABA or glutamate were separately estimated from amongst those astrocytes that had low baseline propagation and those that had high baseline propagation, as described above in *Determining responding cells based on static and propagative events*.

Similarly for each neighboring astrocyte with baseline propagative events, we computed the rate of all events within the baseline period. Separately for GABA and glutamate, we used the baseline event rate across all neighboring astrocytes to define the threshold baseline event rate, *r*_50_, as the 50^th^ percentile of all observed values; cells with baseline rates < *r*_50_ were said to have “low overall baseline event rates”, while cells with fractions ≥ *r*_50_ were said to have “high overall baseline event rates” (Extended Data Fig. 5e, bottom). The fraction of astrocytes that were responders with propagative events to GABA or glutamate were separately estimated from amongst those astrocytes that had low overall baseline event rates and those that had high overall baseline event rates, as above.

To determine the distribution of these responder fractions (amongst astrocytes with low and high baseline propagation, or amongst astrocytes with low and high overall baseline event rates), we performed the same procedure for estimating these fractions on surrogate data generated by hierarchically bootstrapping Ca^2+^ event data 10,000 times (as above in *Propagation probability*). Standard errors were computed as the standard deviation of these empirical distributions.

For each NT, we next sought to determine whether there were significant differences between the fraction of astrocytes that were responders with propagative events amongst cells within the two groupings (*i*.*e*., low vs. high baseline propagation; low vs. high overall baseline event rate). Separately for GABA and glutamate, for each group comparison, we computed the difference between the two responder fractions, as well as the distribution of this difference under an imposed condition in which there was no systematic difference in uncaging response between astrocytes in the two groups. To do this, we performed the same procedure as above for estimating responder fractions in the specified groups (*e*.*g*., “low baseline propagation” and “high baseline propagation”) as well as the difference between the two, but on surrogate data generated by permuting the group labels; this procedure was repeated 10,000 times. As it was unknown *a priori* which group in either comparison—low or high baseline propagation, or low or high overall baseline event rate—would have a higher fraction of responder cells, a two-sided *p*-value was estimated from these surrogate values, as in equation (2).

## Acknowledgments

The authors are grateful to members of the Poskanzer lab for helpful discussions and technical assistance, especially Alba Peinado, Silvia Pittolo, Charlotte Taylor, and Trisha Vaidyanathan. We thank Sae Yokoyama, Gregory Chin, and Nicole DelRosso for technical support; Xuelong Mi and Guoqiang Yu for analytical support; Zhitong (John) Wang for early analysis work; Felice Dunn, Kevin Bender, and Massimo Scanziani for feedback on the project; and Jennifer Thompson for essential administrative support. This work was supported by funding from the National Institutes of Health (project grants R01NS099254 [K.E.P.] and R01MH121446 [K.E.P.]), NSF CAREER (1942360 [K.E.P.]), NSF Graduate Research Fellowship (1650113 [M.K.C.]), CONICET (R.E.) and UCSF Genentech Fellowship (M.E.R.). This material is based upon work supported by the National Science Foundation Graduate Research Fellowship Program under Grant No. 1650113. Any opinions, findings, and conclusions or recommendations expressed in this material are those of the author(s) and do not necessarily reflect the views of the National Science Foundation.

## Author contributions

M.K.C. and K.E.P. conceptualized the project and designed the experiments. M.K.C. performed all *ex vivo* imaging experiments. K.E.P., M.K.C., and M.C. conceptualized the data analysis, and M.K.C. and M.C. performed data analysis with assistance from C.K. V.T. performed IHC, confocal imaging, and related fixed tissue image analysis. M.E.R. performed *in vivo* 2P experiments and related image analysis. R.E. helped with resources related to photoactivatable compounds. M.K.C. and K.E.P. wrote the initial draft, and edited and revised the paper. M.C. contributed to writing the initial draft of the paper and subsequent versions. K.E.P. supervised the research.

## Additional information

**Extended data** Figs. 1–7 and Tables 1–14 are included below.

**Supplementary information** Videos 1–7 are included.

**Correspondence** should be addressed to Kira Poskanzer.

**Extended Data Fig. 1.**
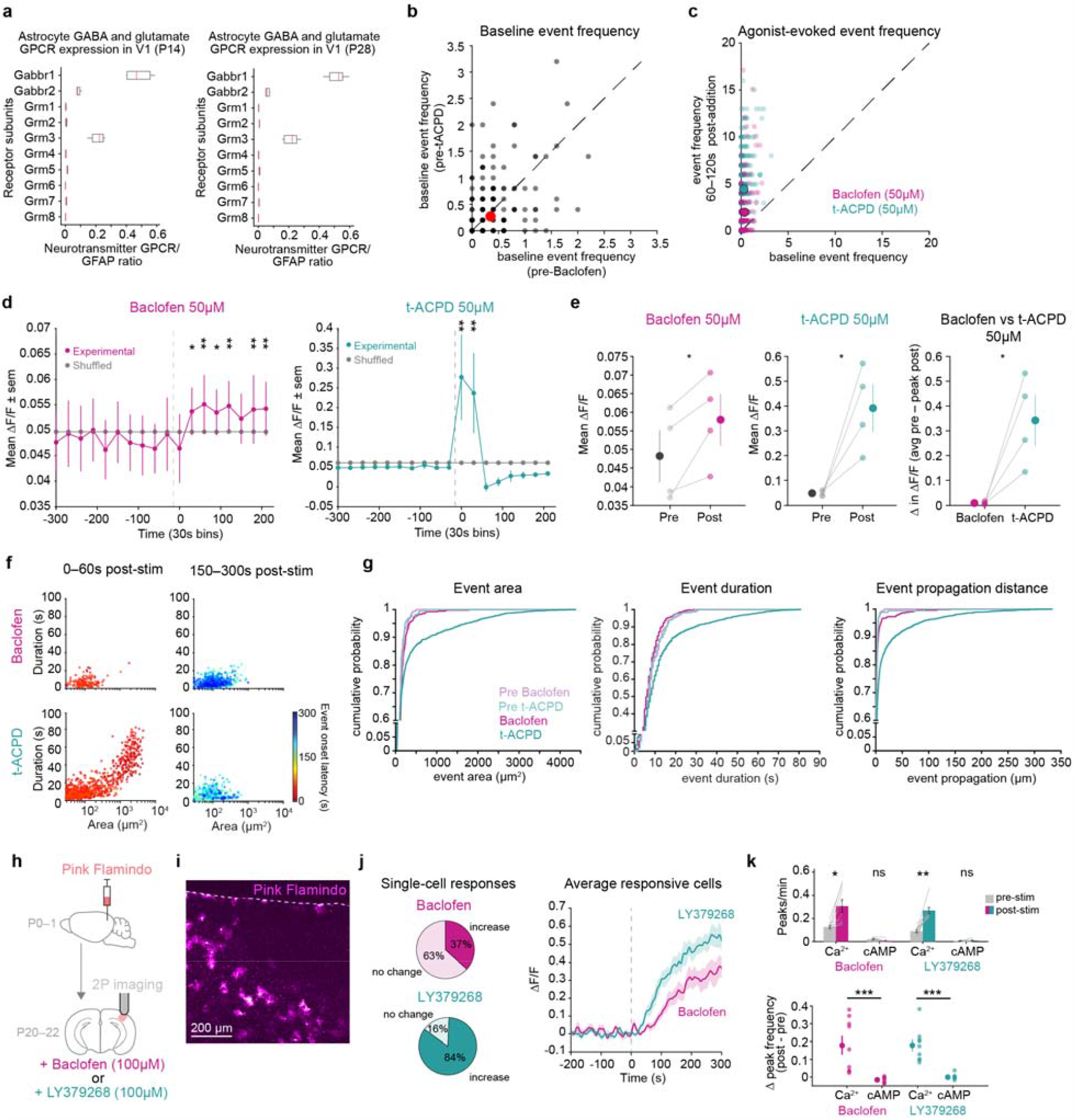
Different responses to activation of astrocytic glutamatergic and GABAergic receptors via pharmacological bath-application. **a**, Ribosomal-mRNA expression in visual cortex astrocytes of P14 (n = 4 biological replicates) and P28 (n = 5 biological replicates) mice from the Farhy-Tselnicker et. al. publicly available dataset (NCBI Gene Expression Omnibus, GSE161398). Visual cortex astrocytes show expression of GABA_B_ receptors and mGluR_3_, but low expression of all other mGluRs, including mGluR_5_ (ref. ^26^). Similar expression levels are found in the Srinivasan et. al. dataset available at http://astrocyternaseq.org/. Ratio of FPKM for the gene of interest / FPKM for GFAP were calculated to normalize for potential differences in the sequencing depth of replicates. **b**, Baseline event frequency for each active astrocyte prior to bath application of baclofen (50μM, x-axis) and t-ACPD (50μM, y-axis). Data shown by astrocyte (grey dots, from n = 4 slices) and mean (red dot). Dashed line = unity line. Baseline event frequencies prior to baclofen and t-ACPD application were compared for each astrocyte using a paired two-sided t-test (*p* = 0.14). **c**, Event frequency for each active astrocyte 300–0s before and 60–120s after addition of agonist (50μM). Data shown by astrocyte (light dots, from n = 4 slices) and mean (solid dots) for baclofen (pink) and t-ACPD (green). Dashed line = unity line; all astrocytes above the unity line display increased activity in presence of agonist. For b & c, 300–0s before addition of agonist was used to calculate mean baseline event frequency (events/60s) per astrocyte; an active astrocyte is any cell with ≥ 1AQuA-detected event. Note the difference in axes between graphs in b & c, reflecting the low baseline event frequency for all astrocytes. **d**, Time-series traces of average ΔF/F in 30 s windows from active cells in each slice. 300–0s before and 0–240s after bath-application of agonist used to calculate event average ΔF/F /30s. Data shown as mean ± sem (n = 4 slices, 4 mice stimulated with 50 μM agonist). Permutation test used to determine significance. *p*-values for all timepoints are in Extended Data Table 3. All traces are aligned to 0s, the frame of agonist entry into the imaging chamber. ΔF/F values were calculated using a moving 10s baseline window, averaging the lower 50% of values in the window. Active cells were cells with ≥ 1 AQuA event detected in either the baclofen or t-ACPD recording. **e**, Left and center: Average ΔF/F before and after bath-application of baclofen (50 μM, left) and t-ACPD (50 μM, center). ΔF/F after bath-application of agonist is from the 30 s time window with the highest average ΔF/F for each slice (“peak post”). Right: Change in average ΔF/F after bath-application of agonist. Data shown as slices (light dots and grey lines, n = 4 slices, 4 mice) and mean ± sem (dark dots and error bars). Paired t-test compares conditions. Baclofen: *p* = 0.046, t-ACPD: *p* = 0.031 and Δ in ΔF/F: *p* = 0.033. **f**, Scatter plots of the area and duration of individual Ca^2+^ events 0–60 s (left) and 150–300 s (right) after bath-application of baclofen (top) or t-ACPD (bottom). Separating events into these two time-windows highlights events occurring early that are covered in Fig. 1d by those with longer onset latencies. Events following bath-application of agonists color-coded by onset time. Dots represent individual Ca^2+^ events from n = 4 slices stimulated with 50μM agonist. Note: these are the same data, with the same onset latency color scale, as shown in Fig. 1d, bottom. **g**, Distributions of event area, duration and propagation 120–0s before (“Pre”) or 0–120s after addition of baclofen (50 μM) or t-ACPD (50 μM). One-way ANOVA followed by Tukey-Kramer Test determine significant pairwise comparisons between conditions. *p-*values for all conditions and features are in Extended Table 4 Note that, for all features, pre-baclofen, pre-tACPD, and baclofen events are not significantly different from one another. Only events following addition of t-ACPD show a rightward shift for all features. **h**, Experimental strategy for Pink Flamindo expression and 2P imaging of astrocytic cAMP in acute cortical slices. **i**, Representative Pink Flamindo fluorescence in V1 FOV; dotted line denotes pia. **j**, Left: Percent of total astrocytes that increase fluorescence or show no change with bath-application of baclofen (top, pink) or mGluR_3_-specific agonist LY379268 (bottom, green) (*n* = 147 astrocytes). Right: Average ∆F/F trace only from responsive cells in each slice (mean ± sem across slices from *n* = 54 responsive astrocytes (baclofen) and 123 responsive astrocytes (LY379268) from *n* = 8 slices, 3 mice). To capture steady-state changes, ∆F/F values were calculated using raw – background fluorescence and a fixed baseline window (frames 1–100), then lowpass filtered at 0.01Hz. **k**, Top: Average Ca^2+^ or cAMP peaks/minute/astrocyte before and after bath-application of baclofen (pink) or LY379268 (green). Data shown as slices (grey lines) and corresponding mean ± sem. Paired t-test compares pre-and post-agonist values for each condition. *p*-values corrected for multiple comparisons using Bonferroni-Holm correction FWER ≤ 0.05. Baclofen: *p* = 0.019 (Ca^2+^) and 0.057 (cAMP). LY379268: *p* = 0.0017 (Ca^2+^) and 0.66 (cAMP). Bottom: Average change in Ca^2+^ or cAMP peaks/minute following bath-application of baclofen (pink) or LY379268 (green). Data shown by slice (light dots) and corresponding mean ± sem (dark dots and error bars). Rank sum tests compare Ca^2+^ and cAMP frequency changes for each agonist. *p* = 0.000082 (baclofen) and 0.000082 (LY379268). Cyto-GCaMP: *n* = 809 active astrocytes (baclofen) and 1033 active astrocytes (LY379268) from *n* = 9 slices, 3 mice. Pink Flamindo: *n* = 147 astrocytes, 8 slices, 3 mice. To detect transient fluctuations, ∆F/F was calculated using a moving 10s baseline window, with peaks determined for each astrocyte if ∆F/F ≥ 3SD above mean baseline ∆F/F.

**Extended Data Fig. 2.**
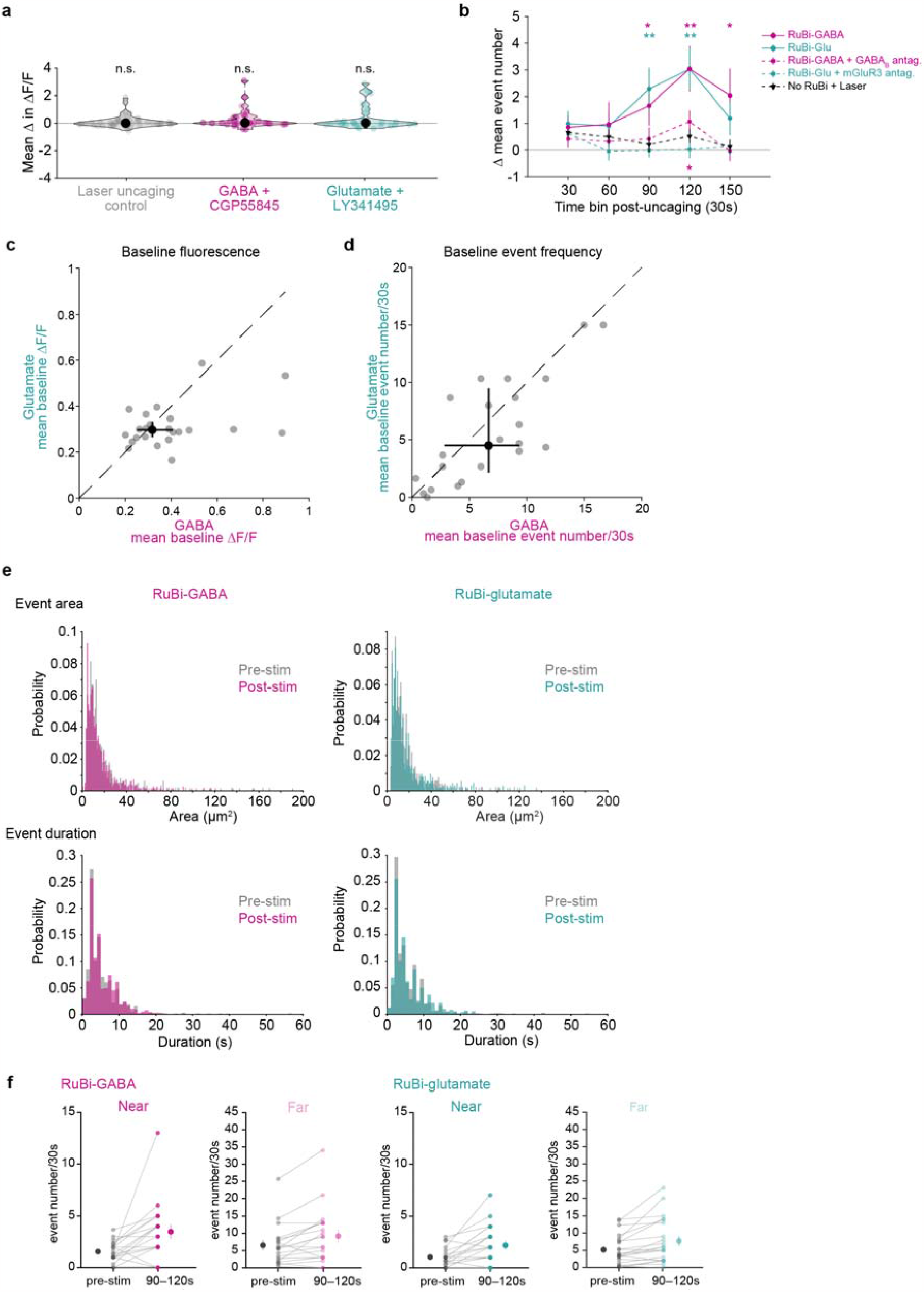
Characterization of, and controls for, increased Ca^2+^ activity in astrocytes directly stimulated by NT uncaging. **a**, Average change in ΔF/F with laser uncaging control (laser stimulation without RuBis, grey, n = 46 astrocytes, 9 slices, 3 mice) and with uncaging in the presence of antagonist (RuBi-GABA + GABA_B_R antagonist [magenta, n = 28 astrocytes, 8 slices, 5 mice] or RuBi-glutamate + mGluR_2/3_ antagonist [green, n = 28 astrocytes, 7 slices, 4 mice]). GABA_B_R antagonized using CGP55845, a potent and selective GABA_B_R antagonist, and mGluR_3_ antagonized using LY341495, a potent mGluR_2/3_ antagonist also known to antagonize other mGluR subtypes at higher concentrations^74^. Data shown by astrocyte, median, 25^th^ and 75^th^ percentile. Wilcoxon signed-rank test compares change from baseline. *p*-values corrected for multiple comparisons using Bonferroni-Holm correction with FWER ≤ 0.05. Laser uncaging control: *p* = 0.50, RuBi-GABA + CGP55845: *p* = 0.11 and RuBi-glutamate + LY341495: *p* = 0.41. **b**, Event frequency change after NT uncaging (GABA: solid magenta lines, n = 27 astrocytes, 7 slices, 4 mice; glutamate: solid green lines, n = 24 astrocytes, 7 slices, 4 mice), NT uncaging in the presence of antagonist (dotted magenta and green lines), and laser uncaging control (dotted black line). 90–0s before and 0–150s after uncaging used to calculate event number/30s. Data shown by mean ± sem. Permutation test used to determine significance. *p*-values for all conditions and timepoints are in Extended Data Table 5. **c**, Baseline fluorescence prior to GABA and glutamate uncaging. 90–0s before uncaging used to calculate mean ∆F/F per cell. Data shown by cell (grey dots, n = 24 astrocytes), median, and 25^th^ and 75^th^ percentile (black dots and crosshairs). Dashed line = unity line. Wilcoxon signed-rank test shows no significant difference between baseline fluorescence of directly stimulated astrocytes prior to GABA and glutamate uncaging (*p* = 0.089). **d**, Baseline event frequency prior to GABA and glutamate uncaging. 90–0s before uncaging used to calculate mean number of events/30s for each cell. Data shown by cell (grey dots, n = 24 astrocytes), median and 25^th^ and 75^th^ percentile (black dots and crosshairs). Dashed line = unity line. Wilcoxon signed-rank test shows no significant difference between baseline event frequency of directly stimulated astrocytes prior to GABA and glutamate uncaging (*p* = 0.068). **e**, Distribution of event area and duration pre-mand post-uncaging of RuBi-GABA (left) and RuBi-glutamate (right) from “responder” uncaging cells. Detected events 120 s pre- and post-uncaging are included from n = 19 astrocytes, 7 slice, 4 mice (GABA) and n = 21 astrocytes, 7 slices, 4 mice (glutamate). Rank-sum test compares pre- and post-uncaging event features. Area: *p* = 0.58 (GABA) and 0.95 (glutamate). Duration: *p* = 0.083 (GABA) and 0.13 (glutamate). **f**, Event frequency in responding astrocytes directly stimulated with NT. Events from directly stimulated astrocytes were separated into events near and far from GABA and glutamate uncaging. 90–0s before used to calculate average event number/30s (“pre-stim”). Data shown by cell (light dots and grey lines) and mean ± sem (dark dots and error bars).

**Extended Data Fig. 3.**
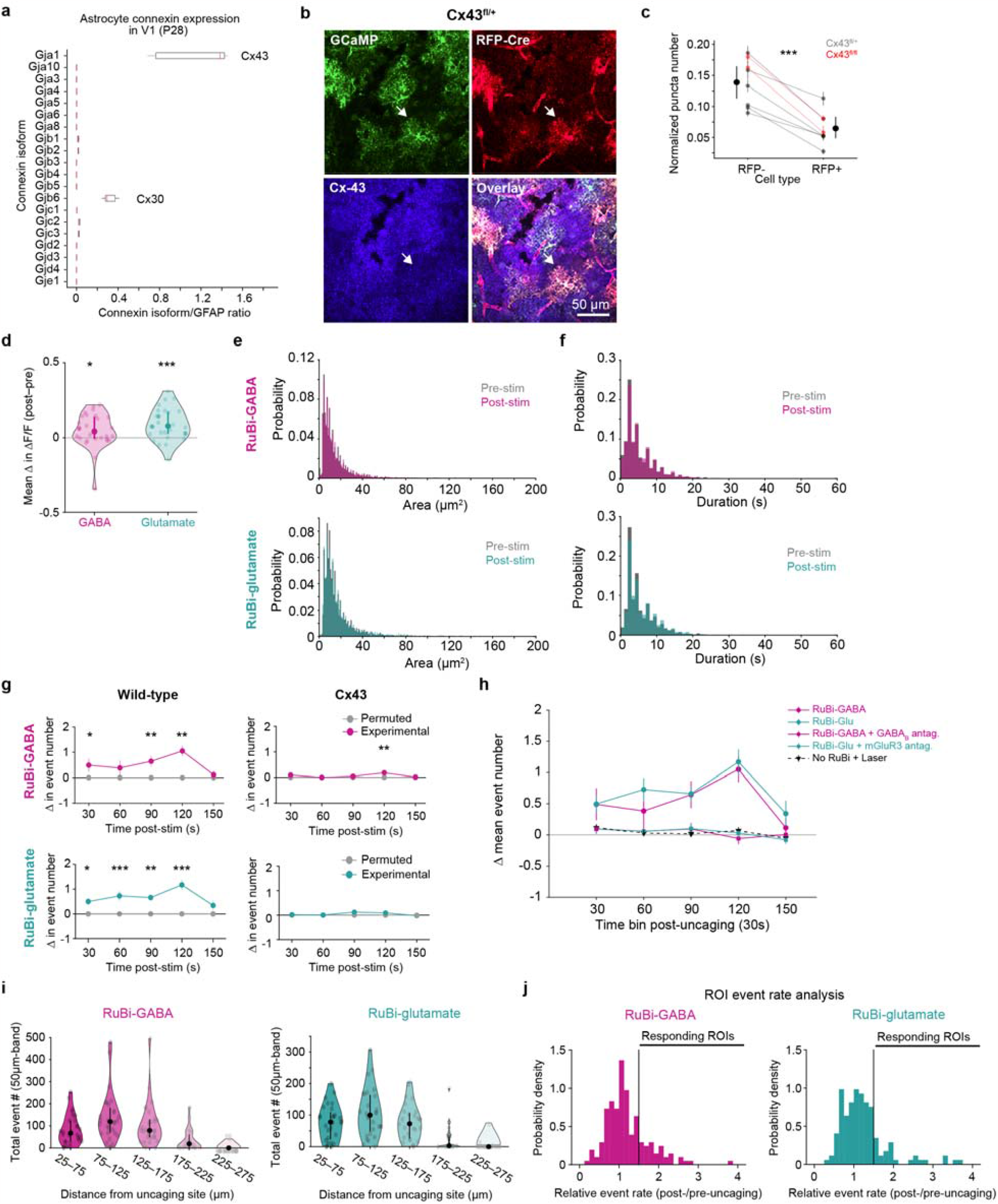
Confirmation of Cx43 knockdown and network-level controls after NT uncaging. **a**, Ribosomal-mRNA expression in visual cortex astrocytes of P28 mice (n = 5 biological replicates) from the Farhy-Tselnicker et. al. publicly available dataset (NCBI Gene Expression Omnibus, GSE161398). Visual cortex astrocytes preferentially express Cx43 (Gja1) over other connexins, including Cx30 (Gjb6). Similar expression levels are found in the Srinivasan et. al. dataset available at http://astrocyternaseq.org. Ratio of FPKM for the gene of interest / FPKM for GFAP were calculated to normalize for potential differences in the sequencing depth of replicates. **b**, Representative micrographs of immunohistochemistry in a Cx43^fl/+^ slice demonstrating reduced numbers of Cx43 puncta in Cre^+^ astrocytes. White arrow points to individual cell expressing GCaMP (green) and RFP-Cre (red), with reduced Cx43 (blue). **c**, Average Cx43 puncta/astrocyte in RFP-Cre^-^ and RFP-Cre^+^ astrocytes; puncta counts are normalized by area of each astrocyte. Data are shown by mouse averages (light dots, error bars and connecting lines, grey = Cx43^fl/+^ and red = Cx43^fl/fl^ mice) and mean ± sem (dark dots and error bars). Cx43 puncta counts were similar for Cx43^fl/+^ and Cx43^fl/fl^ mice; data from both genotypes were pooled together for all analyses and referred to as Cx43^floxed^. Paired two-sided t-test compares average Cx43 puncta counts in RFP-Cre^-^ and RFP-Cre^+^ astrocytes. *p* = 0.00013. **d**, Average change in ∆F/F in WT astrocyte networks after RuBi-GABA (magenta) and RuBi-glutamate (green) uncaging. Data shown by trial/FOV, median and 25^th^ and 75^th^ percentile. (For d–i, WT: n = 28 networks, 7 slices, 4 mice). Wilcoxon signed-rank test compares change from baseline. *p* = 0.016 (GABA) and 0.00032 (glutamate). **e**,**f**, Distribution of event area and duration pre- and post-uncaging of RuBi-GABA (top) and RuBi-glutamate (bottom). Detected events 120 s pre- and post-uncaging are included. Rank-sum test compares pre- and post-uncaging event features. Area: *p* = 0.025 (GABA) and 0.0050 (glutamate). Duration: *p* = 0.063 (GABA) and 0.0000045 (glutamate). **g**,**h**, Event frequency change in neighboring astrocytes after GABA (g, top) and glutamate (g, bottom) uncaging in WT and Cx43^floxed^ slices (n = 61 networks, 16 slices, 8 mice). WT data from g replotted in h (circular markers) with laser uncaging control (laser stimulation without RuBis, dotted black line and triangular markers, n = 48 networks, 9 slices, 3 mice) and with uncaging in the presence of antagonist (RuBi-GABA + GABA_B_R antagonist [magenta line and square markers, n = 32 networks, 8 slices, 5 mice] or RuBi-glutamate + mGluR_2/3_ antagonist [green line and square markers, n = 28 networks, 7 slices, 4 mice]). 90–0s before and 0–150s after uncaging used to calculate event number/30s in neighboring astrocytes with ≥ 1 AQuA-detected event. Data shown by mean ± sem. Permutation test used to determine significance. *p*-values for all conditions and timepoints are in Extended Data Table 7. **i**, Total number of AQuA-detected events in 50μm bands radiating out from the uncaging site. All events 90s before and 150s after NT uncaging are included. Data shown by trial/FOV, median and 25^th^ and 75^th^ percentile. **j**, Distribution of relative event rates from 20×20μm ROIs following uncaging of RuBi-GABA (left) and RuBi-glutamate (right). Validation for threshold used to define ROIs with increased activity post-uncaging; chosen threshold: ≥ 50% event frequency increase post-uncaging.

**Extended Data Fig. 4.**
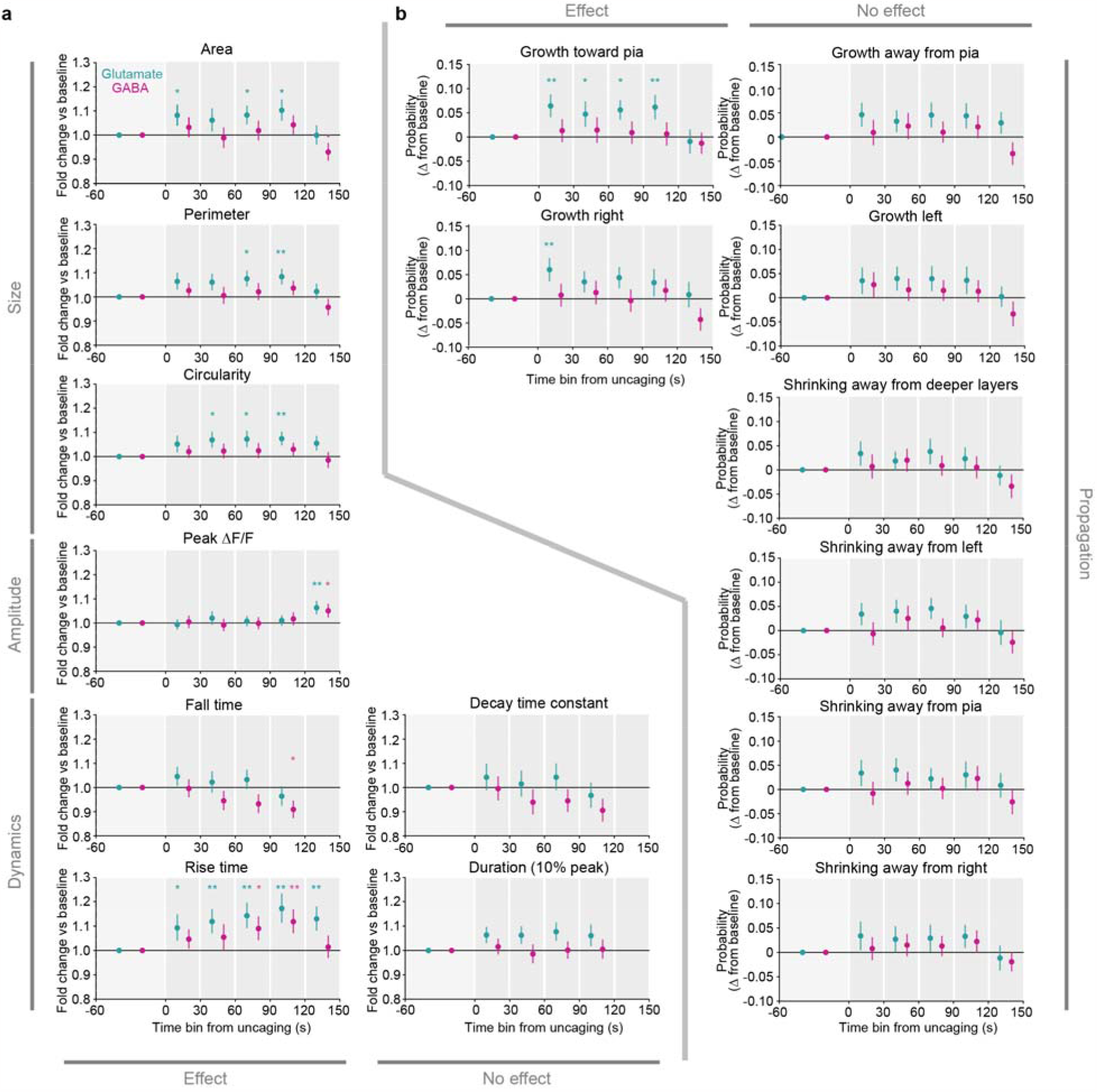
Change in individual astrocyte Ca^2+^ event features post NT-uncaging. **a**, Fold change in indicated Ca^2+^ event features among all events from all neighboring cells after GABA or glutamate uncaging, relative to 60–0s pre-uncaging. Data shown as overall fold change ± standard error determined from hierarchical bootstrapping (Methods; GABA: *n* = 142 cells in 28 FOV; glutamate: *n* = 120 cells in 27 FOV). Two-sided *p*- and *q*-values for changes versus baseline were obtained by circularly shifting each cell’s events in time (see Methods; Extended Data Table 12). **b**, Change in the probability of a Ca^2+^ event growing or shrinking in the indicated direction among all events from neighboring cells after GABA or glutamate uncaging, relative to 60–0s pre-uncaging. Data shown as overall probability ± standard error determined from hierarchical bootstrapping (Methods; GABA: *n* = 142 cells in 28 FOV; glutamate: *n* = 120 cells in 27 FOV). Two-sided *p*- and *q*-values for changes versus baseline were obtained by circularly shifting each cell’s events in time (see Methods; Extended Data Table 13).

**Extended Data Fig. 5.**
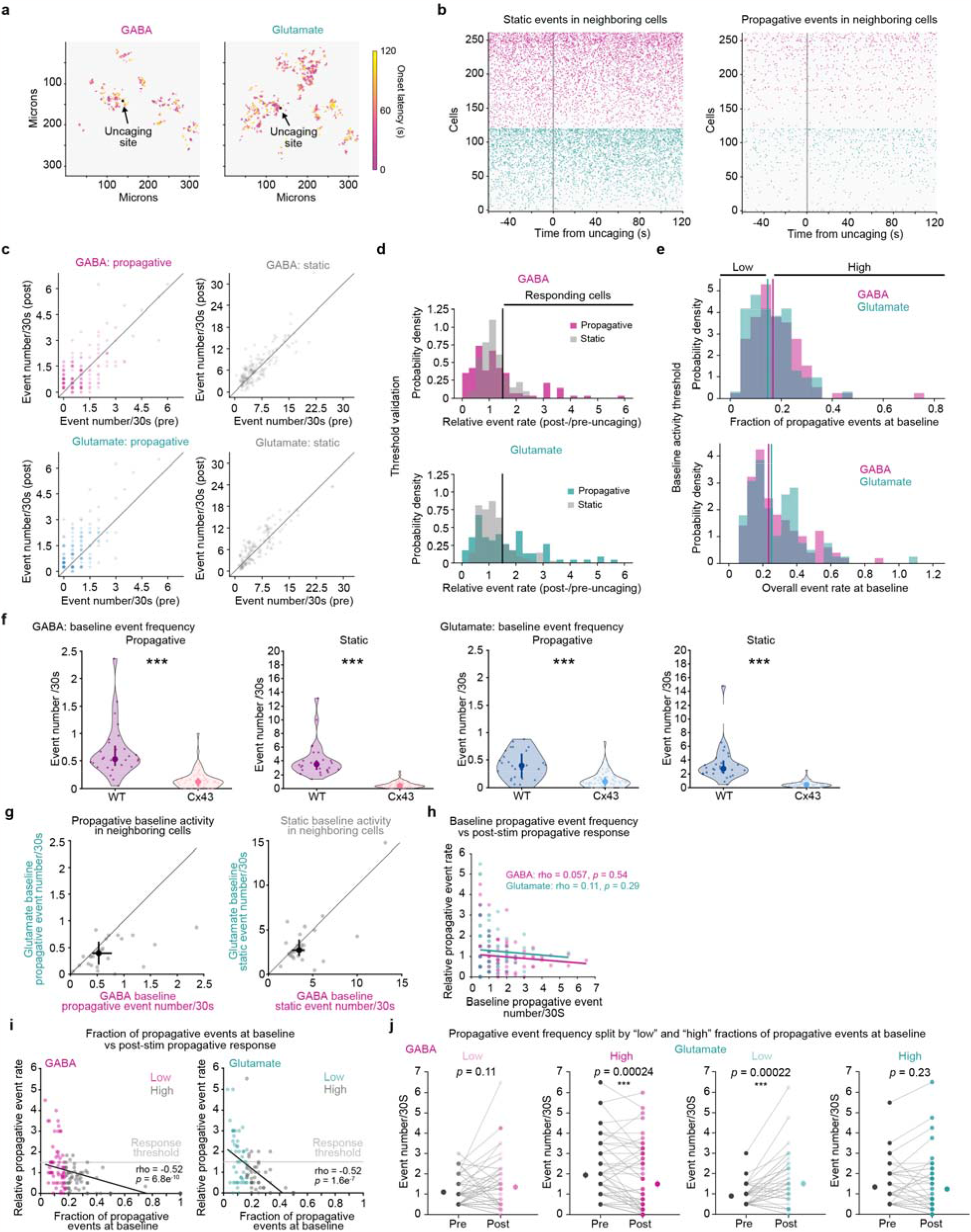
Validating changes in propagative event activity following NT-uncaging. **a**, Representative spatial maps of Ca^2+^ events in the same astrocyte network 0–120 s after either GABA (left) or glutamate (right) uncaging. Events are color-coded by onset time. Black dot = NT uncaging site. Events from all time-points are distributed throughout the imaging field. There is no visible wavefront of activity traveling across the imaging field or emanating from the uncaging site. Note that all panels except for (f) are data from WT slices. **b**, Raster plots of Ca^2+^ event onsets for static (left) or propagative (right) events before and after GABA (magenta) or glutamate (green) uncaging. Raster plots show all neighboring cells (astrocytes not directly stimulated by NT-uncaging) from all FOVs, with each row showing events from an individual astrocyte. Within each NT and event type, cells were sorted by the overall rate of static events from 0–120s post-uncaging (*i*.*e*., the same sorting was used for the left and right raster plots). Grey line = NT uncaging start. **c**, Scatter plots of event rates (event number/30s) within neighboring cells during the period 60–0s pre-uncaging (*x*-axis) versus 0–120s post-uncaging (*y*-axis). Rates of propagative (left) and static (right) events are shown for recordings of GABA (top) and glutamate (bottom) uncaging. Dots are individual neighboring cells; darker dots indicate multiple overlapping cells. **d**, Distribution of post-/pre-uncaging ratio of static (grey) or propagative (color) event rates among neighboring cells with any baseline events of the corresponding type, after GABA (magenta, top) or glutamate (green, bottom) uncaging. Ratios computed per-cell as the rate from 0–120s post-uncaging divided by the rate from 60–0s pre-uncaging. Vertical black lines indicate the threshold used to determine “responding” cells in Fig. 4h, l, m and Extended Data Fig. 6e (*i*.*e*., ≥ 1.5-fold). **e**, Top: Distribution of the fraction of events during the baseline window (60–0s pre-uncaging) that were propagative in each neighboring cell before GABA (magenta) or glutamate (green) uncaging, among those cells that had any baseline propagative activity. Vertical magenta and green lines indicate the thresholds (50^th^ percentile) for recordings of GABA and glutamate uncaging, respectively, used in Fig. 4m left to delineate “Low” and “High” fraction of propagative events at baseline among neighboring cells. Bottom: Distribution of the overall event rate during the baseline window of 60–0s pre– GABA (magenta) or glutamate (green) uncaging, in each neighboring cell that had baseline propagative activity. Vertical magenta and green lines indicate the thresholds (50^th^ percentile) for recordings of GABA and glutamate uncaging, respectively, used in Fig. 4m right to delineate cells with “Low” and “High” overall event rates at baseline. **f**, Baseline propagative (left) and static (right) event frequencies of astrocytes in WT or Cx43^floxed^ slices. Baseline period: 90–0s prior to uncaging. Individual data points show average event rate from active neighboring astrocytes (≥ 1 AQuA-detected event during recording) for each FOV. Data shown by FOV (WT: *n* = 28 FOV for GABA and glutamate, 7 slices, 4 mice; Cx43^floxed^: *n* = 63 FOV for GABA and 61 FOV for glutamate, 16 slices, 8 mice), median, 25^th^ and 75^th^ percentile. Wilcoxon rank sum test compares WT and Cx43^floxed^ baseline event frequencies (GABA: *p* = 1.6e-10 [propagative], 7.7e-14 [static]; glutamate: *p* = 9.0e-7 [propagative], 1.1e-12 [static]). **g**, Baseline propagative (left) and static (right) event frequencies in WT networks prior to GABA and glutamate uncaging. 90–0s before uncaging used to calculate mean number of events/30s. Event rate per FOV calculated by averaging the event rates of active astrocytes in the FOV (≥ 1 AQuA-detected event during the recording), excluding the uncaging astrocyte. Data shown by FOV (grey dots, n = 28), median, 25^th^ and 75^th^ percentile (black dot and crosshairs). Wilcoxon signed-rank test compares baseline event frequencies prior to GABA and glutamate uncaging (*p* = 0.00022 [propagative] and 0.052 [static]). **h**, Spearman correlation between baseline propagative event rate and relative post-stim propagative event rate for neighboring cells in GABA (magenta) and glutamate (turquoise) recordings. Data shown by individual neighboring astrocyte (for h–i, n = 121 cells [GABA], 91 cells [glutamate] with ≥ 1 baseline propagative event). For h–i, 60–0s before uncaging used for baseline window and relative post-stim propagative rate calculated as in d. **i**, Spearman correlation between fraction of propagative events at baseline and relative post-stim propagative event rate for neighboring cells in GABA (left) and glutamate (right) recordings. Data shown by individual neighboring astrocyte color-coded by baseline activity composition category (“low” in magenta or turquoise, “high” in grey). Light grey horizontal line = response threshold (responders ≥ 1.5-fold increase in propagative activity from baseline). Note a majority of astrocytes responding to either NT (at or above the response threshold line) display a low fraction of propagative events at baseline. **j**, Propagative event frequency pre- and post-uncaging for neighboring cells with “low” and “high” fractions of propagative events at baseline (as for Fig. 4m, left). 60–0s before (“Pre”) and 0–120s after (“Post”) used to calculate average event number/30s. Data shown by cell (light dots and grey lines; n = 61 cells [GABA “low”], 60 cells [GABA “high”], 46 cells [glutamate “low”], 45 cells [glutamate “high”]) and mean ± sem (dark dots and error bars). Wilcoxon signed-rank test compare pre-and post-stim frequencies for each category.

**Extended Data Fig. 6.**
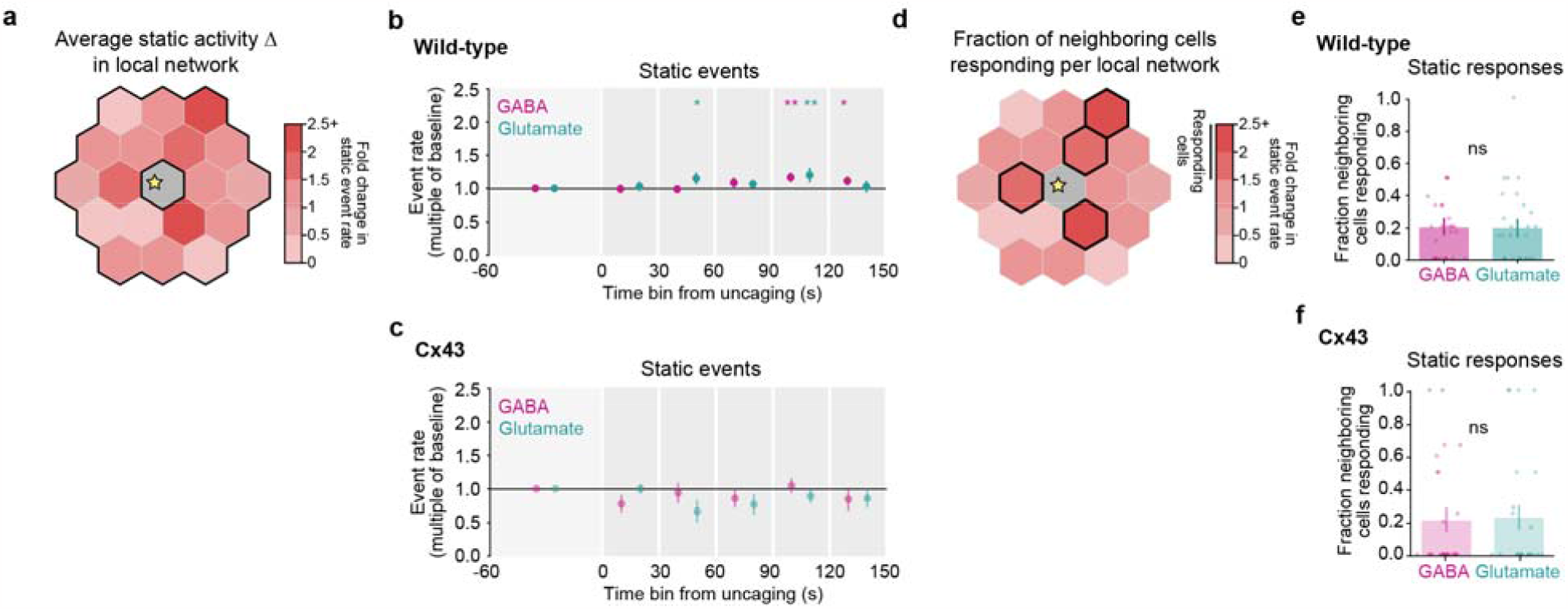
Static activity changes in the local astrocyte network are similar in response to GABA and glutamate. **a**, Analysis schematic illustrating average static activity change across all neighboring cells in the local network, as reported in b and c. Heterogeneous responses of individual neighboring cells are averaged in b and c. **b–c**, Fold-change in rate of static Ca^2+^ events among neighboring cells after GABA or glutamate uncaging in acute slices from WT mice (**b**) or Cx43^floxed^ mice (**c**), relative to 60–0s pre-uncaging. Data shown as median across FOVs ± standard error via hierarchical bootstrapping (Methods; *n* for all conditions are in Extended Data Table 9). One-sided *p*- and *q*-values were obtained via circular permutation testing (Methods; Extended Data Table 10); *: *q* < 0.05, **: *q* < 0.01. **d**, Analysis schematic illustrating the fraction of neighboring cells per FOV that respond to NT with increases in static activity, as reported in e and f. Note that a subset of neighboring cells in the local network exhibit activity changes following NT uncaging. **e–f**, Fraction of neighboring cells per FOV with ≥ 50% increase in static Ca^2+^ events (*responding*) after GABA or glutamate uncaging in WT (**e**) or Cx43^floxed^ slices (**f**). Data shown as mean ± sem via hierarchical bootstrapping; dots denote individual FOVs (see Methods; *n* for all conditions are in Extended Data Table 9). Permutation testing was used to compare fraction of cells responding to GABA and glutamate in WT slices (*p* = 1.0) and Cx43^floxed^ slices (static: *p* = 1.0).

**Extended Data Fig. 7.**
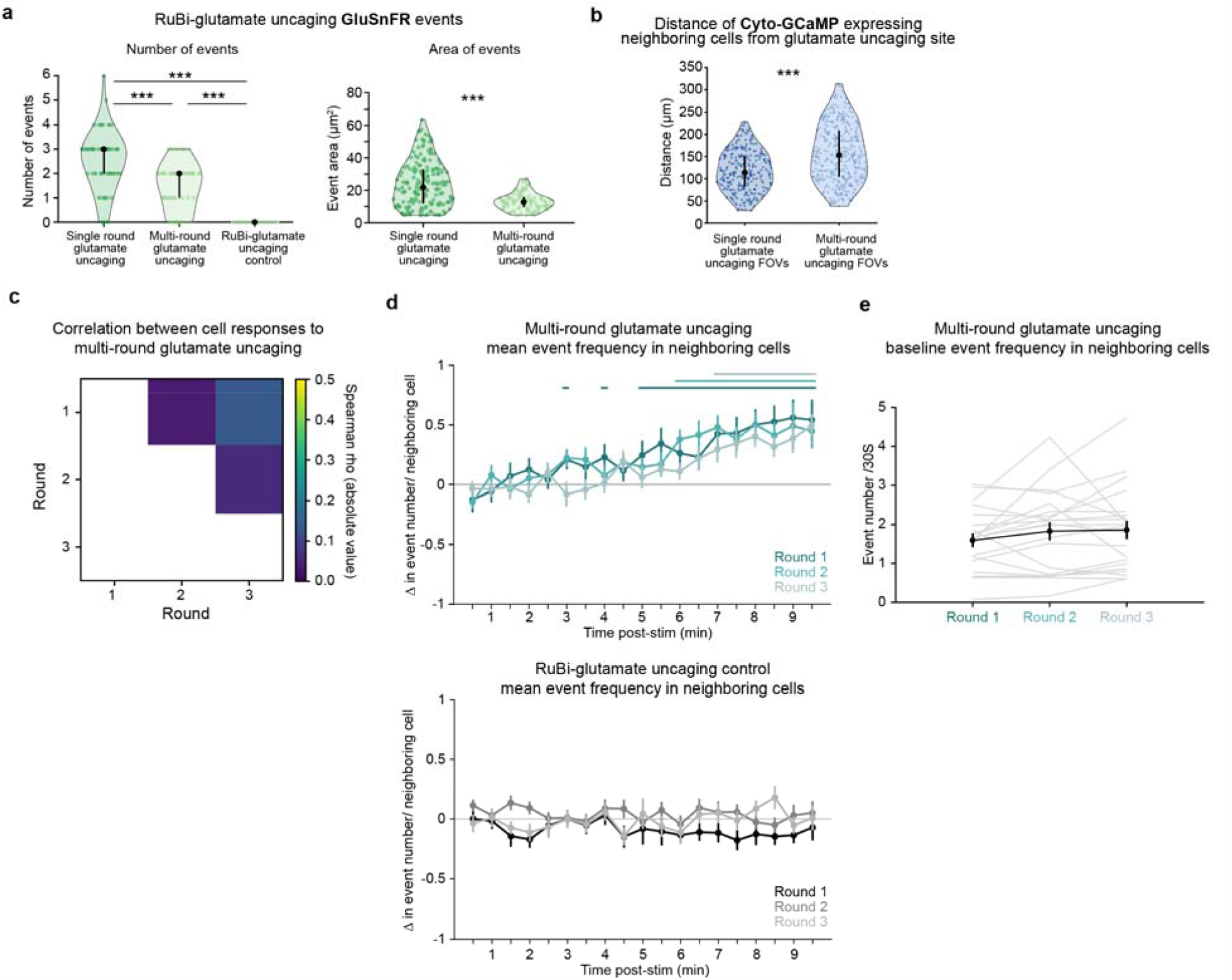
Individual neighboring astrocytes exhibit variable Ca^2+^ responses across multiple rounds of glutamate uncaging. **a**, GluSnFR event features after RuBi-glutamate uncaging for three types of uncaging datasets. For number of events/uncaging site (left), data shown by uncaging trial, median, 25^th^ and 75^th^ percentile. For GluSnFR event area (right), data shown by GluSnFR event, median, 25^th^ and 75^th^ percentile (single round glutamate uncaging: *n* = 72 trials, 12 recordings, 4 slices, 2 mice; multi-round glutamate uncaging: *n* = 66 trials, 11 recordings, 2 slices, 1 mouse; RuBi-glutamate uncaging control: *n* = 66 trials, 11 recordings, 2 slices, 1 mouse). For number of events, one-way ANOVA followed by Tukey-Kramer Test determine significant pairwise comparisons between laser stimulation conditions. *p* = 9.7e-10 (single round glutamate uncaging v multi-round glutamate uncaging), 9.6e-10 (single round glutamate uncaging v RuBi-glutamate uncaging control) and 9.6e-10 (multi-round glutamate uncaging vs RuBi-glutamate uncaging control). For event area, rank sum test compares single round glutamate uncaging vs. multi-round glutamate uncaging, *p* = 3.6e-10. All datasets were collected in the presence of RuBi-glutamate. For single round and multi-round glutamate uncaging, the uncaging laser power was set to 70 A.U. (∼8mW at the sample). Laser re-alignment between these datasets leads to a small difference in amount of glutamate uncaged with laser stimulation (see event area on right). For RuBi-glutamate uncaging controls, the uncaging laser power was set to 25 A.U. (∼2mW at the sample), a stimulation that did not lead to detectable glutamate uncaging (see event number on left). **b**, Distance of Cyto-GCaMP-expressing neighboring astrocytes from the glutamate uncaging site. Distance measured from the centroid of each neighboring astrocyte to the centroid of the uncaging site. Data shown by active astrocyte (≥ 1 AQuA-detected event 0–300s from recording onset), median, 25^th^ and 75^th^ percentile (single round glutamate uncaging: *n* = 28 FOV, 7 slices, 4 mice; multi-round glutamate uncaging: *n* = 23 FOV, 9 slices, 5 mice). Rank sum test compares datasets; *p* = 3.4e-15. **c**, Correlation between the propagative Ca^2+^ responses of individual neighboring cells to multiple rounds of glutamate uncaging. Individual cells’ binary responses to glutamate uncaging are not significantly correlated across rounds (Spearman rho = 0.040, *p* = 1.0, n = 32 cells, 15 recordings, 8 slices, 5 mice [round 1 vs 2]; Spearman rho = 0.14, *p* = 0.70, n = 30 cells, 16 recordings, 7 slices, 5 mice [round 1 vs 3]; Spearman rho = 0.059, *p* = 0.74, n = 38 cells, 17 recordings, 8 slices, 5 mice [round 2 vs 3]), showing that the response of an individual cell is variable from round to round. In each round, activity was recorded 150–0s before and 0–600s following uncaging, with glutamate uncaged over an area of ∼12μm^2^ (as in a, right “Multi-round glutamate uncaging”). Rounds of imaging/uncaging for each FOV were separated by ≥ 25 minutes. Cells included in analysis for each round had ≥ 1 propagative event during 60–0s before uncaging. Responding cells exhibited ≥ 50% increase in propagative event frequency 300–420s following uncaging, a time window in which activity began to increase across rounds, compared to 60–0s before uncaging. **d**, Event frequency change in neighboring astrocytes across three rounds of glutamate uncaging (top) and RuBi-glutamate uncaging controls (bottom). 90–0s before and 0–570s after uncaging used to calculate mean event number/30s in active astrocytes (astrocytes in the local network with ≥1 AQuA-detected event during recording, excluding the stimulated cell). Data shown by mean ± sem (multi-round glutamate uncaging: n = 23 FOV for Round 1 and 3, 21 FOV for Round 2, 9 slices, 5 mice; RuBi-glutamate uncaging control: n = 20 FOV, 8 slices, 5 mice). Permutation test used to determine significance. *p*-values for all conditions, rounds, and time points are in Extended Data Table 14. The responses in multi-round glutamate uncaging are delayed compared to the single round glutamate uncaging dataset (Extended Data Fig. 3g). Two factors may account for this delay. First, less NT is released in the multi-round glutamate uncaging dataset (a). Second, the distance of astrocytes in the local network from the uncaging site is greater in the multi-round uncaging dataset compared to the single round uncaging dataset (b). **e**, Baseline event frequencies for neighboring astrocytes across three rounds of glutamate uncaging. 90–0s before uncaging used to calculate mean event number/30s/active astrocytes in each FOV. Data shown FOV (light grey lines, n = 21 FOV, 9 slices, 5 mice) and mean ± sem (black dots and error bars). Repeated measures ANOVA compares baseline frequencies across rounds (F(2,40) = 1.51, *p* = 0.23).

**Extended Data Table 1.** Statistics for Fig. 1c: Change in event frequency in astrocytes during bath-application of agonist. Permutation testing used to identify time-points with changes in event frequency greater than chance for each agonist. *p*-values corrected for multiple comparisons using Benjamini-Yekutieli procedure with FDR ≤ 0.05. Adjusted *p*-values < 0.05 are bold.

| **Condition** | **Time relative to agonist entry (s)** | ***p*-values (raw)** | ***p*-values (adjusted)** |
| --- | --- | --- | --- |
| **Baclofen** | **-240 – -180** | 0.598040195980402 | 1.90331632015370 |
|  | **-180 – -120** | 0.429157084291571 | 1.82110765113965 |
|  | **-120 – -60** | 0.374462553744626 | 1.82110765113965 |
|  | **-60 – 0** | 0.570242975702430 | 1.90331632015370 |
|  | **0–60** | 0.339466053394661 | 1.82110765113965 |
|  | **60–120** | 9.99900009999000e-05 | **0.000848605615628913** |
|  | **120–180** | 9.99900009999000e-05 | **0.000848605615628913** |
|  | **180–240** | 9.99900009999000e-05 | **0.000848605615628913** |
| **t-ACPD** | **-240 – -180** | 0.950704929507049 | 2.80096426865250 |
|  | **-180 – -120** | 0.939206079392061 | 2.80096426865250 |
|  | **-120 – -60** | 0.851514848515149 | 2.80096426865250 |
|  | **-60 – 0** | 0.990100989901010 | 2.80096426865250 |
|  | **0–60** | 9.99900009999000e-05 | **0.000848605615628913** |
|  | **60–120** | 9.99900009999000e-05 | **0.000848605615628913** |
|  | **120–180** | 9.99900009999000e-05 | **0.000848605615628913** |
|  | **180–240** | 0.0241975802419758 | 0.154021919236648 |

**Extended Data Table 2.** Statistics for Fig. 1e–h: Dose-response curves. Paired t-tests at each concentration compare response to each agonist for each feature. *p*-values corrected for multiple comparisons using Bonferroni-Holm correction with FWER ≤ 0.05. Adjusted *p*-values < 0.05 are bold.

| **Feature** | **Agonist concentration ($\mu\text{M}$)** | ***p*-values (raw)** | ***p*-values (adjusted)** |
| --- | --- | --- | --- |
| **Percent field active** | 5 | 0.106884262919877 | 0.106884262919877 |
|  | 25 | 0.00453636186896923 | **0.00907272373793845** |
|  | 50 | 0.000175165474580620 | **0.000700661898322481** |
|  | 100 | 0.000644835771994738 | **0.00193450731598421** |
| **Event area** | 5 | 0.134291909278406 | 0.134291909278406 |
|  | 25 | 0.00336246922231183 | **0.0100874076669355** |
|  | 50 | 0.00274661524985296 | **0.0109864609994119** |
|  | 100 | 0.0355015432722231 | 0.0710030865444463 |
| **Event duration** | 5 | 0.794523834669017 | 0.794523834669017 |
|  | 25 | 0.00693357100768626 | **0.0208007130230588** |
|  | 50 | 0.000707221042301198 | **0.00282888416920479** |
|  | 100 | 0.0222444305319271 | **0.0444888610638542** |
| **Event propagation** | 5 | 0.503037904758348 | 0.503037904758348 |
|  | 25 | 0.00201639415469164 | **0.00806557661876658** |
|  | 50 | 0.00332572662050501 | **0.00997717986151502** |
|  | 100 | 0.0102174453241840 | **0.0204348906483680** |

**Extended Data Table 3.**
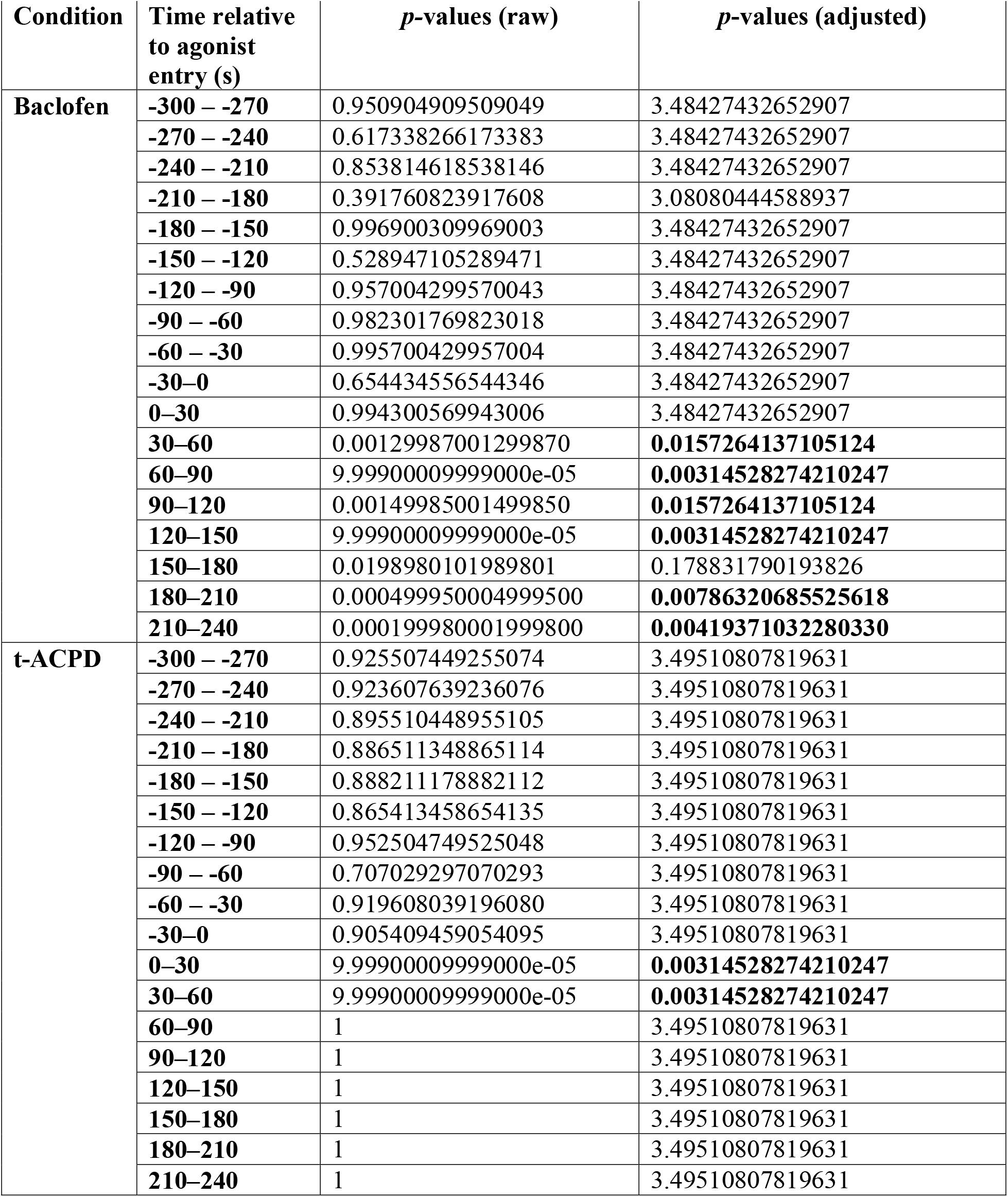
Statistics for Extended Data Fig. 1d: Mean ∆F/F in astrocytes during bath-application of agonist. Permutation testing used to identify time-points with increases in ∆F/F greater than chance for each agonist. *p*-values corrected for multiple comparisons using Benjamini-Yekutieli procedure with FDR ≤ 0.05. Adjusted *p*-values < 0.05 are bold.

**Extended Data Table 4.** Statistics for Extended Data Fig. 1g: Event features pre- and post-agonist addition. Comparison of distributions of event area, duration, and propagation 120–0s before (“Pre”) or 0– 120s after (“Post”) addition of baclofen (50 μM) or t-ACPD (50 μM). One-way ANOVA followed by Tukey-Kramer Test determine significant pairwise comparisons between conditions. *p*-values < 0.05 are bold, greyed-out cells are pairwise comparisons that are not relevant.

| Feature | Test | Comparison | *p*-values |
| --- | --- | --- | --- |
| **Area** | 1-way ANOVA |  | **2.64502036644004e-13** |
|  | Tukey-Kramer Test | Bac_pre v Bac_post | 0.898277274752255 |
|  |  | Bac_pre v tACPD_pre | 0.995109946477354 |
|  |  | Bac_pre v tACPD_post | 7.59031018511269e-05 |
|  |  | Bac_post v tACPD_pre | 0.975557200328527 |
|  |  | Bac_post v tACPD_post | **8.83569961640518e-09** |
|  |  | tACPD_pre v tACPD_post | **0.000158328341554381** |
| **Duration** | 1-way ANOVA |  | **2.90200679997044e-19** |
|  | Tukey-Kramer Test | Bac_pre v Bac_post | 0.705179159520994 |
|  |  | Bac_pre v tACPD_pre | 0.997414358965489 |
|  |  | Bac_pre v tACPD_post | 0.000778858034344410 |
|  |  | Bac_post v tACPD_pre | 0.822905967024531 |
|  |  | Bac_post v tACPD_post | **3.76825814996096e-09** |
|  |  | tACPD_pre v tACPD_post | **0.000133775981740025** |
| **Propagation distance** | 1-way ANOVA |  | **5.06378633615887e-15** |
|  | Tukey-Kramer Test | Bac_pre v Bac_post | 0.913414390535003 |
|  |  | Bac_pre v tACPD_pre | 0.998952459006666 |
|  |  | Bac_pre v tACPD_post | 3.10437553621146e-05 |
|  |  | Bac_post v tACPD_pre | 0.959725706670736 |
|  |  | Bac_post v tACPD_post | **4.12490908097141e-09** |
|  |  | tACPD_pre v tACPD_post | **3.44219605847851e-05** |
| **ΔF/F** | 1-way ANOVA |  | **1.35022940218368e-148** |
|  | Tukey-Kramer Test | Bac_pre v Bac_post | 0.787076385709762 |
|  |  | Bac_pre v tACPD_pre | 0.988259466617291 |
|  |  | Bac_pre v tACPD_post | 3.76825814996096e-09 |
|  |  | Bac_post v tACPD_pre | 0.518577476864315 |
|  |  | Bac_post v tACPD_post | **3.76825814996096e-09** |
|  |  | tACPD_pre v tACPD_post | **3.76825814996096e-09** |

**Extended Data Table 5.** Statistics for Extended Data Fig. 2b: Change in event frequency in astrocytes directly stimulated with NT. Permutation testing used to identify time-points with changes in event frequency greater than chance for each condition. *p*-values corrected for multiple comparisons using Benjamini-Yekutieli procedure with FDR ≤ 0.05. Adjusted *p*-values < 0.05 are bold.

| **Condition** | **Time relative to uncaging (s)** | ***p*-values (raw)** | ***p*-values (adjusted)** |
| --- | --- | --- | --- |
| **GABA** | 0–30 | 0.196880311968803 | 0.449543378995434 |
|  | 30–60 | 0.149385061493851 | 0.426369863013699 |
|  | 60–90 | 0.0115988401159884 | **0.0441400304414003** |
|  | 90–120 | 9.99900009999000e-05 | **0.00114155251141553** |
|  | 120–150 | 0.00229977002299770 | **0.0131278538812785** |
| **Glutamate** | 0–30 | 0.118788121187881 | 0.339041095890411 |
|  | 30–60 | 0.153984601539846 | 0.351598173515982 |
|  | 60–90 | 0.000399960003999600 | **0.00228310502283105** |
|  | 90–120 | 9.99900009999000e-05 | **0.00114155251141553** |
|  | 120–150 | 0.0591940805919408 | 0.225266362252664 |
| **GABA + CGP55845** | 0–30 | 0.237476252374763 | 0.906392694063927 |
|  | 30–60 | 0.363263673632637 | 1.03681506849315 |
|  | 60–90 | 0.238176182381762 | 0.906392694063927 |
|  | 90–120 | 0.00279972002799720 | **0.0319634703196347** |
|  | 120–150 | 0.927407259274073 | 2.11757990867580 |
| **Glutamate + LY341495** | 0–30 | 0.0266973302669733 | 0.304794520547945 |
|  | 30–60 | 0.883111688831117 | 2.23515981735160 |
|  | 60–90 | 0.978902109789021 | 2.23515981735160 |
|  | 90–120 | 0.919108089191081 | 2.23515981735160 |
|  | 120–150 | 0.650334966503350 | 2.23515981735160 |
| **Laser uncaging control** | 0–30 | 0.0105989401059894 | 0.121004566210046 |
|  | 30–60 | 0.0492950704929507 | 0.187595129375951 |
|  | 60–90 | 0.407059294070593 | 1.16181506849315 |
|  | 90–120 | 0.0379962003799620 | 0.187595129375951 |
|  | 120–150 | 0.622337766223378 | 1.42100456621005 |

**Extended Data Table 6.** Statistics for Fig. 3h: Change in event frequency in the astrocyte network via Sholl-like analysis. Permutation testing used to identify time-points with changes in event frequency greater than chance for each distance band and NT. *p*-values corrected for multiple comparisons using Benjamini-Yekutieli procedure with FDR ≤ 0.05. Adjusted *p*-values < 0.05 are bold.

| NT | Distance from uncaging ($\mu\text{m}$) | Time relative to uncaging (s) | *p*-values (raw) | *p*-values (adjusted) |
| --- | --- | --- | --- | --- |
| GABA | 25–75 | 0–30 | 0.0569943005699430 | 0.216894977168950 |
|  |  | 30–60 | 0.798720127987201 | 1.82374429223744 |
|  |  | 60–90 | 0.0429957004299570 | 0.216894977168950 |
|  |  | 90–120 | 0.00599940005999400 | 0.0684931506849315 |
|  |  | 120–150 | 0.535546445355465 | 1.52853881278539 |
|  | 75–125 | 0–30 | 0.0666933306669333 | 0.276636225266362 |
|  |  | 30–60 | 0.187281271872813 | 0.534531963470320 |
|  |  | 60–90 | 0.0726927307269273 | 0.276636225266362 |
|  |  | 90–120 | 0.00299970002999700 | **0.0342465753424658** |
|  |  | 120–150 | 0.307269273072693 | 0.701598173515982 |
|  | 125–175 | 0–30 | 0.0418958104189581 | 0.0956621004566210 |
|  |  | 30–60 | 0.0140985901409859 | 0.0536529680365297 |
|  |  | 60–90 | 0.0192980701929807 | 0.0550799086757991 |
|  |  | 90–120 | 9.99900009999000e-05 | **0.00114155251141553** |
|  |  | 120–150 | 0.00379962003799620 | **0.0216894977168950** |
| glutamate | 25–75 | 0–30 | 0.00169983001699830 | **0.00646879756468798** |
|  |  | 30–60 | 0.00119988001199880 | **0.00646879756468798** |
|  |  | 60–90 | 0.00799920007999200 | **0.0228310502283105** |
|  |  | 90–120 | 9.99900009999000e-05 | **0.00114155251141553** |
|  |  | 120–150 | 0.0128987101289871 | **0.0294520547945206** |
|  | 75–125 | 0–30 | 0.255074492550745 | 0.728025114155251 |
|  |  | 30–60 | 0.00459954004599540 | **0.0262557077625571** |
|  |  | 60–90 | 0.00779922007799220 | **0.0296803652968037** |
|  |  | 90–120 | 9.99900009999000e-05 | **0.00114155251141553** |
|  |  | 120–150 | 0.576442355764424 | 1.31621004566210 |
|  | 125–175 | 0–30 | 0.113988601139886 | 0.260273972602740 |
|  |  | 30–60 | 0.0355964403559644 | 0.135464231354642 |
|  |  | 60–90 | 0.00949905009499050 | 0.0542237442922375 |
|  |  | 90–120 | 9.99900009999000e-05 | **0.00114155251141553** |
|  |  | 120–150 | 0.0554944505549445 | 0.158390410958904 |

**Extended Data Table 7.** Statistics for Extended Data Fig. 3g,h: Change in event frequency in individual astrocytes in the network. Permutation testing used to identify time-points with changes in event frequency greater than chance for each condition. *p*-values corrected for multiple comparisons using Benjamini-Yekutieli procedure with FDR ≤ 0.05. Adjusted *p*-values < 0.05 are bold.

| Condition | Time relative to uncaging (s) | *p*-values (raw) | *p*-values (adjusted) |
| --- | --- | --- | --- |
| GABA WT | 0–30 | 0.00409959004099590 | **0.0156012176560122** |
|  | 30–60 | 0.0294970502949705 | 0.0841894977168950 |
|  | 60–90 | 0.000299970002999700 | **0.00171232876712329** |
|  | 90–120 | 9.99900009999000e-05 | **0.00114155251141553** |
|  | 120–150 | 0.519848015198480 | 1.18698630136986 |
| Glutamate WT | 0–30 | 0.00479952004799520 | **0.0136986301369863** |
|  | 30–60 | 9.99900009999000e-05 | **0.000570776255707763** |
|  | 60–90 | 0.000299970002999700 | **0.00114155251141553** |
|  | 90–120 | 9.99900009999000e-05 | **0.000570776255707763** |
|  | 120–150 | 0.0480951904809519 | 0.109817351598174 |
| GABA Cx43<sup>flox</sup> | 0–30 | 0.0586941305869413 | 0.335045662100457 |
|  | 30–60 | 0.943105689431057 | 2.15342465753425 |
|  | 60–90 | 0.373262673732627 | 1.42047184170472 |
|  | 90–120 | 0.000499950004999500 | **0.00570776255707763** |
|  | 120–150 | 0.814718528147185 | 2.15342465753425 |
| Glutamate Cx43<sup>flox</sup> | 0–30 | 0.668933106689331 | 1.90924657534247 |
|  | 30–60 | 0.878712128787121 | 2.00639269406393 |
|  | 60–90 | 0.0154984501549845 | 0.176940639269406 |
|  | 90–120 | 0.0617938206179382 | 0.352739726027397 |
|  | 120–150 | 0.666933306669333 | 1.90924657534247 |
| GABA + CGP55845 | 0–30 | 0.229877012298770 | 1.32363013698630 |
|  | 30–60 | 0.460253974602540 | 1.32363013698630 |
|  | 60–90 | 0.234976502349765 | 1.32363013698630 |
|  | 90–120 | 0.463753624637536 | 1.32363013698630 |
|  | 120–150 | 0.971902809719028 | 2.21917808219178 |
| Glutamate + LY341495 | 0–30 | 0.181181881811819 | 0.997716894977169 |
|  | 30–60 | 0.427057294270573 | 1.21889269406393 |
|  | 60–90 | 0.160683931606839 | 0.997716894977169 |
|  | 90–120 | 0.660733926607339 | 1.50867579908676 |
|  | 120–150 | 0.262173782621738 | 0.997716894977169 |
| Laser uncaging control | 0–30 | 0.00899910008999100 | 0.102739726027397 |
|  | 30–60 | 0.498050194980502 | 1.42151826484018 |
|  | 60–90 | 0.668633136686331 | 1.52671232876712 |
|  | 90–120 | 0.130786921307869 | 0.746575342465753 |
|  | 120–150 | 0.239876012398760 | 0.912861491628615 |

**Extended Data Table 8.**
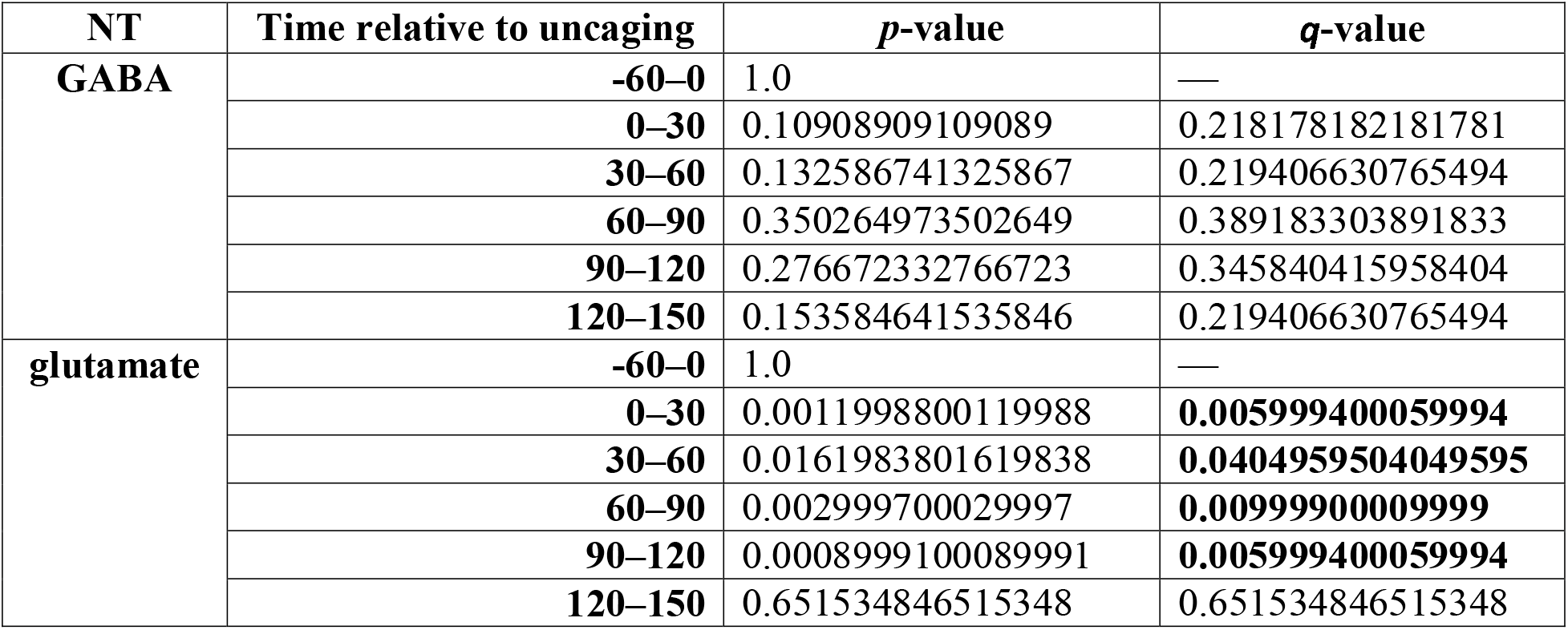
Statistics for Fig. 4b: Change in probability of Ca^2+^ events propagating toward or away from pia compared to baseline. Two-sided permutation testing was used to identify time bins with changes in propagative event probability compared to baseline. *p*-values were adjusted across tested time bins using the Benjamini-Hochberg procedure to obtain *q*-values. *q*-values < 0.05 are bold.

| **NT** | **Time relative to uncaging** | ***p*-value** | ***q*-value** |
| --- | --- | --- | --- |
| **GABA** | **-60–0** | 1.0 | — |
|  | **0–30** | 0.10908909109089 | 0.218178182181781 |
|  | **30–60** | 0.132586741325867 | 0.219406630765494 |
|  | **60–90** | 0.350264973502649 | 0.389183303891833 |
|  | **90–120** | 0.276672332766723 | 0.345840415958404 |
|  | **120–150** | 0.153584641535846 | 0.219406630765494 |
| **glutamate** | **-60–0** | 1.0 | — |
|  | **0–30** | 0.0011998800119988 | **0.005999400059994** |
|  | **30–60** | 0.0161983801619838 | **0.0404959504049595** |
|  | **60–90** | 0.002999700029997 | **0.00999900009999** |
|  | **90–120** | 0.0008999100089991 | **0.005999400059994** |
|  | **120–150** | 0.651534846515348 | 0.651534846515348 |

**Extended Data Table 9.** N for Fig. 4f, h, j, k and Extended Data Fig. 6: Fold-change in rate of static or propagative Ca^2+^ events in neighboring cells post NT-uncaging in WT and Cx43^floxed^ mice. Event rate changes were used to calculate the fraction of neighboring cells/FOV responding to NT-uncaging for each condition (Fig. 4 h and k and Extended Data Fig. 6 e–f).

| Genotype | Event Category | NT | N |  |  |  |  |
| --- | --- | --- | --- | --- | --- | --- | --- |
|  |  |  | event | cell | FOV | slice | mice |
| WT | static | GABA | 8417 | 142 | 28 | 7 | 4 |
|  |  | glutamate | 6998 | 120 | 27 | 7 | 4 |
|  | propagative | GABA | 1358 | 135 | 28 | 7 | 4 |
|  |  | glutamate | 1112 | 115 | 27 | 7 | 4 |
| Cx43<sup>flox</sup> | static | GABA | 1566 | 60 | 28 | 14 | 8 |
|  |  | glutamate | 1215 | 47 | 23 | 14 | 8 |
|  | propagative | GABA | 487 | 57 | 28 | 14 | 8 |
|  |  | glutamate | 348 | 47 | 23 | 14 | 8 |

**Extended Data Table 10.** Statistics for Fig. 4f, j and Extended Data Fig. 6 b–c: Fold-change in rate of static or propagative Ca^2+^ events among neighboring cells post NT-uncaging in WT and Cx43^floxed^ mice. One-sided permutation test used to identify time bins with static or propagative event rate increases compared to baseline. *p*-values were adjusted across tested time bins using the Benjamini-Hochberg procedure to obtain *q*-values. *q*-values < 0.05 are bold.

| Genotype | Event Category | NT | Time relative to uncaging | *p*-value | *q*-value |
| --- | --- | --- | --- | --- | --- |
| WT | static | GABA | -60–0 | 1.0 | — |
|  |  |  | 0–30 | 0.5649435056494350 | 0.5888411158884110 |
|  |  |  | 30–60 | 0.5888411158884110 | 0.5888411158884110 |
|  |  |  | 60–90 | 0.0560943905609439 | 0.1121887811218870 |
|  |  |  | 90–120 | 0.0012998700129987 | **0.0064993500649935** |
|  |  |  | 120–150 | 0.0150984901509849 | **0.0377462253774622** |
|  |  | glutamate | -60–0 | 1.0 | — |
|  |  |  | 0–30 | 0.2686731326867310 | 0.3735876412358760 |
|  |  |  | 30–60 | 0.0066993300669933 | **0.0223311002233110** |
|  |  |  | 60–90 | 0.1374862513748620 | 0.2291437522914370 |
|  |  |  | 90–120 | 0.0009999000099990 | **0.0064993500649935** |
|  |  |  | 120–150 | 0.2988701129887010 | 0.3735876412358760 |
|  | propagative | GABA | -60–0 | 1.0 | — |
|  |  |  | 0–30 | 0.0341965803419658 | 0.0683931606839316 |
|  |  |  | 30–60 | 0.6876312368763120 | 0.7640347076403470 |
|  |  |  | 60–90 | 0.2723727627237270 | 0.3404659534046590 |
|  |  |  | 90–120 | 0.0726927307269273 | 0.1211545512115450 |
|  |  |  | 120–150 | 0.9524047595240470 | 0.9524047595240470 |
|  |  | glutamate | -60–0 | 1.0 | — |
|  |  |  | 0–30 | 0.0071992800719928 | **0.0179982001799820** |
|  |  |  | 30–60 | 0.0071992800719928 | **0.0179982001799820** |
|  |  |  | 60–90 | 0.0000999900009999 | **0.0004999500049995** |
|  |  |  | 90–120 | 0.0000999900009999 | **0.0004999500049995** |
|  |  |  | 120–150 | 0.2249775022497750 | 0.3213964317853920 |
| Cx43<sup>flox</sup> | static | GABA | -60–0 | 1.0 | — |
|  |  |  | 0–30 | 0.961603839616038 | 0.995600439956004 |
|  |  |  | 30–60 | 0.698030196980302 | 0.995600439956004 |
|  |  |  | 60–90 | 0.859614038596140 | 0.995600439956004 |
|  |  |  | 90–120 | 0.198480151984801 | 0.995600439956004 |
|  |  |  | 120–150 | 0.874012598740125 | 0.995600439956004 |
|  |  | glutamate | -60–0 | 1.0 | — |
|  |  |  | 0–30 | 0.559344065593440 | 0.995600439956004 |
|  |  |  | 30–60 | 0.995600439956004 | 0.995600439956004 |
|  |  |  | 60–90 | 0.915308469153084 | 0.995600439956004 |
|  |  |  | 90–120 | 0.700929907009299 | 0.995600439956004 |
|  |  |  | 120–150 | 0.798820117988201 | 0.995600439956004 |
|  | propagative | GABA | -60–0 | 1.0 | — |
|  |  |  | 0–30 | 0.812918708129187 | 0.933684409336844 |
|  |  |  | 30–60 | 0.939206079392060 | 0.939206079392060 |
|  |  |  | 60–90 | 0.444755524447555 | 0.933684409336844 |
|  |  |  | 90–120 | 0.830316968303169 | 0.933684409336844 |
|  |  |  | 120–150 | 0.813818618138186 | 0.933684409336844 |
|  |  | glutamate | -60–0 | 1.0 | — |
|  |  |  | 0–30 | 0.793020697930207 | 0.933684409336844 |
|  |  |  | 30–60 | 0.834616538346165 | 0.933684409336844 |
|  |  |  | 60–90 | 0.397960203979602 | 0.933684409336844 |
|  |  |  | **90–120** | 0.379562043795620 | 0.933684409336844 |
|  |  |  | **120–150** | 0.840315968403159 | 0.933684409336844 |

**Extended Data Table 11.** N for Fig. 4m: Fraction of neighboring cells responding to NT-uncaging with propagative event frequency increases separated by baseline activity levels. “All” (static and propagative) events were used to calculate overall baseline event rate and fraction of propagative events in the baseline period to separate cells into “low” and “high” overall baseline activity or “low” and “high” baseline propagation, respectively. Propagative events were used to categorize cells as “responders” or “non-responders” to NT-uncaging.

| Event Category | NT | N |  |  |  |  |
| --- | --- | --- | --- | --- | --- | --- |
|  |  | event | cell | FOV | slice | mice |
| all | GABA | 9775 | 142 | 28 | 7 | 4 |
|  | glutamate | 8110 | 120 | 27 | 7 | 4 |
| propagative | GABA | 1358 | 135 | 28 | 7 | 4 |
|  | glutamate | 1112 | 115 | 27 | 7 | 4 |

**Extended Data Table 12.**
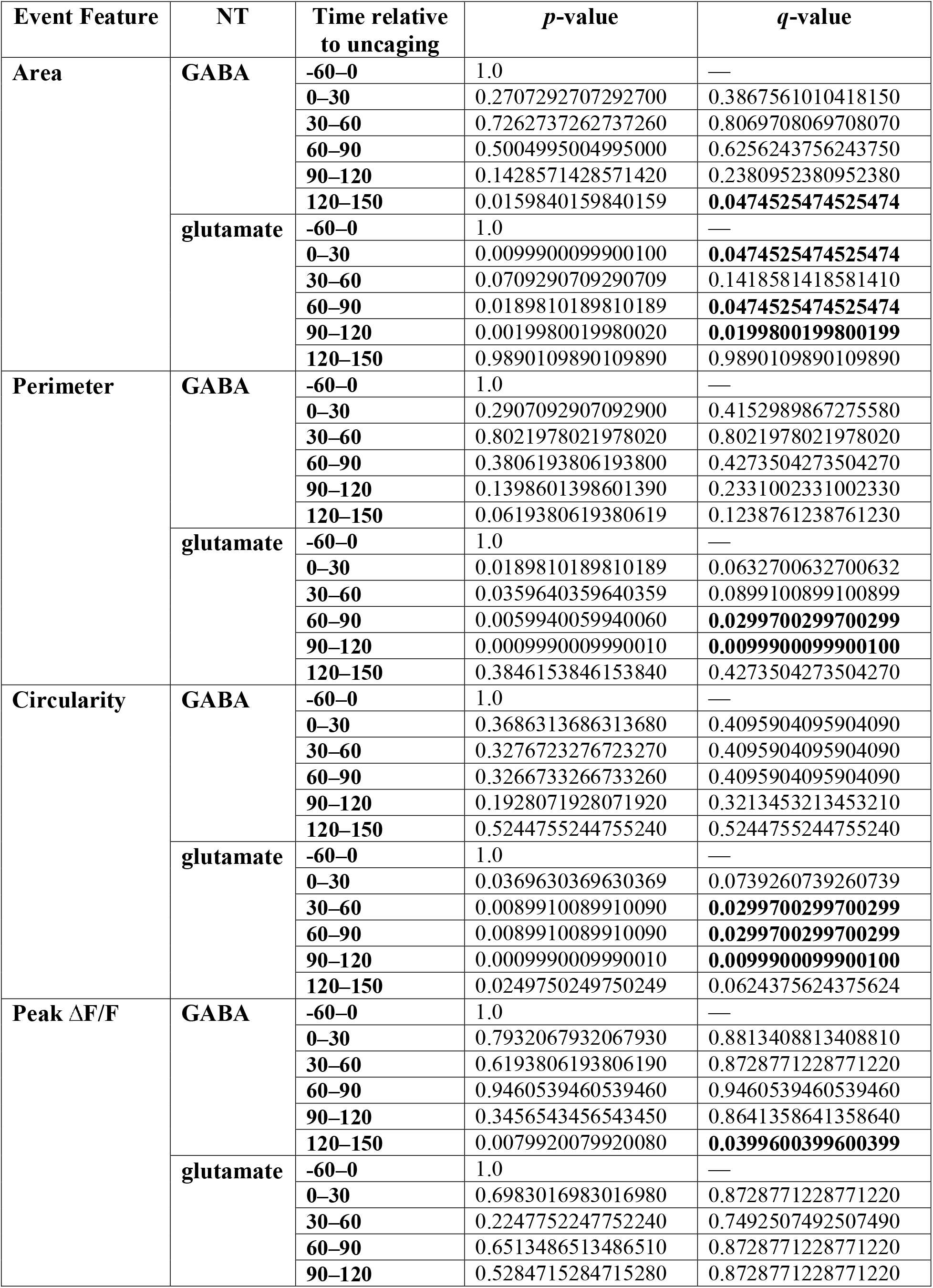
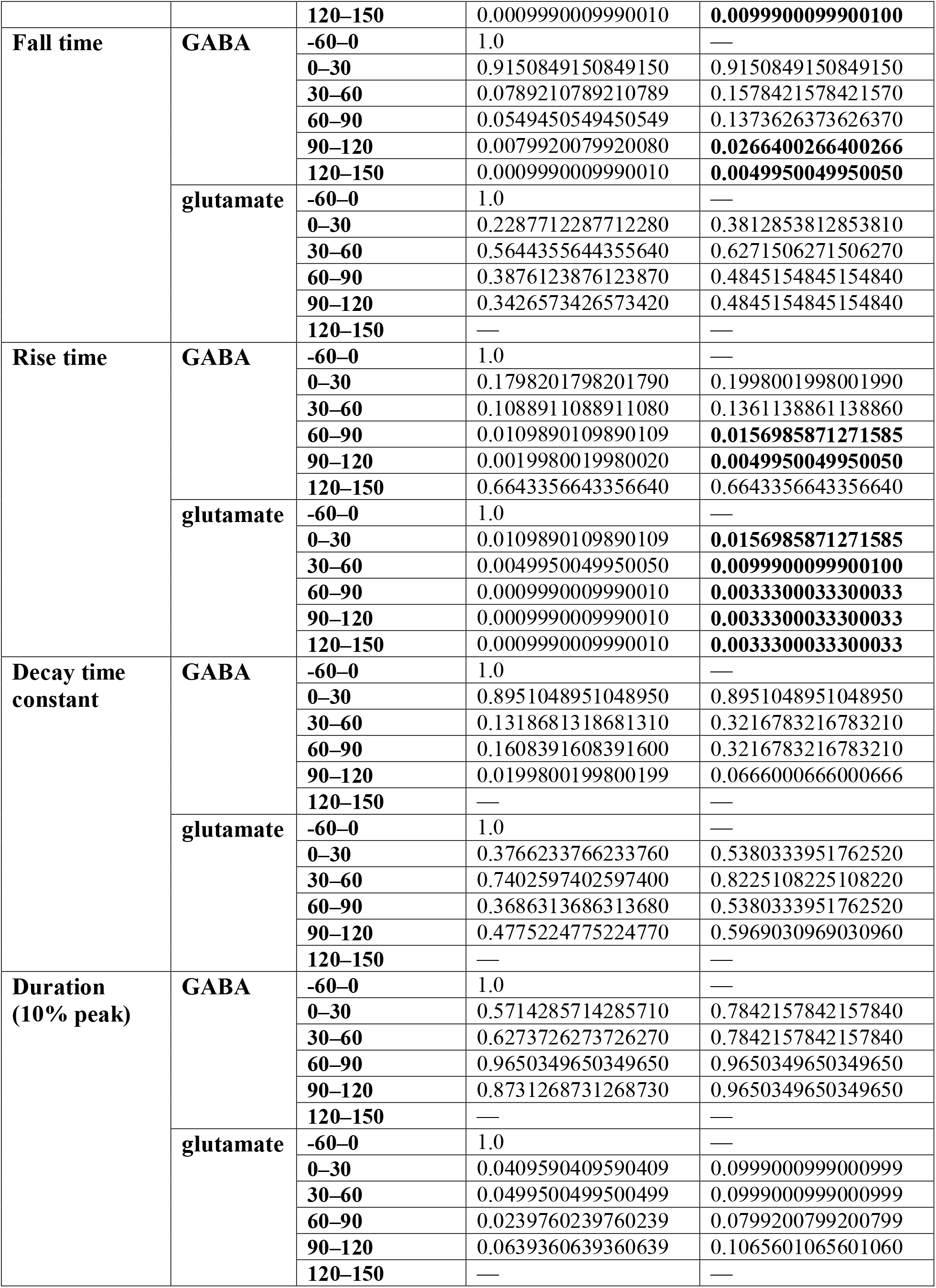
Statistics for Extended Data Fig. 4a: Fold change in individual Ca^2+^ events features compared to baseline. Two-sided permutation testing was used to identify time bins with changes in event features compared to baseline. *p*-values were adjusted across tested time bins using the Benjamini-Hochberg procedure to obtain *q*-values. *q*-values < 0.05 are bold.

| Event Feature | NT | Time relative to uncaging | *p*-value | *q*-value |
| --- | --- | --- | --- | --- |
| Area | GABA | -60–0 | 1.0 | — |
|  |  | 0–30 | 0.2707292707292700 | 0.3867561010418150 |
|  |  | 30–60 | 0.7262737262737260 | 0.8069708069708070 |
|  |  | 60–90 | 0.5004995004995000 | 0.6256243756243750 |
|  |  | 90–120 | 0.1428571428571420 | 0.2380952380952380 |
|  |  | 120–150 | 0.0159840159840159 | **0.0474525474525474** |
|  | glutamate | -60–0 | 1.0 | — |
|  |  | 0–30 | 0.0099900099900100 | **0.0474525474525474** |
|  |  | 30–60 | 0.0709290709290709 | 0.1418581418581410 |
|  |  | 60–90 | 0.0189810189810189 | **0.0474525474525474** |
|  |  | 90–120 | 0.0019980019980020 | **0.0199800199800199** |
|  |  | 120–150 | 0.9890109890109890 | 0.9890109890109890 |
| Perimeter | GABA | -60–0 | 1.0 | — |
|  |  | 0–30 | 0.2907092907092900 | 0.4152989867275580 |
|  |  | 30–60 | 0.8021978021978020 | 0.8021978021978020 |
|  |  | 60–90 | 0.3806193806193800 | 0.4273504273504270 |
|  |  | 90–120 | 0.1398601398601390 | 0.2331002331002330 |
|  |  | 120–150 | 0.0619380619380619 | 0.1238761238761230 |
|  | glutamate | -60–0 | 1.0 | — |
|  |  | 0–30 | 0.0189810189810189 | 0.0632700632700632 |
|  |  | 30–60 | 0.0359640359640359 | 0.0899100899100899 |
|  |  | 60–90 | 0.0059940059940060 | **0.0299700299700299** |
|  |  | 90–120 | 0.0009990009990010 | **0.0099900099900100** |
|  |  | 120–150 | 0.3846153846153840 | 0.4273504273504270 |
| Circularity | GABA | -60–0 | 1.0 | — |
|  |  | 0–30 | 0.3686313686313680 | 0.4095904095904090 |
|  |  | 30–60 | 0.3276723276723270 | 0.4095904095904090 |
|  |  | 60–90 | 0.3266733266733260 | 0.4095904095904090 |
|  |  | 90–120 | 0.1928071928071920 | 0.3213453213453210 |
|  |  | 120–150 | 0.5244755244755240 | 0.5244755244755240 |
|  | glutamate | -60–0 | 1.0 | — |
|  |  | 0–30 | 0.0369630369630369 | 0.0739260739260739 |
|  |  | 30–60 | 0.0089910089910090 | **0.0299700299700299** |
|  |  | 60–90 | 0.0089910089910090 | **0.0299700299700299** |
|  |  | 90–120 | 0.0009990009990010 | **0.0099900099900100** |
|  |  | 120–150 | 0.0249750249750249 | 0.0624375624375624 |
| Peak $\Delta F/F$ | GABA | -60–0 | 1.0 | — |
|  |  | 0–30 | 0.7932067932067930 | 0.8813408813408810 |
|  |  | 30–60 | 0.6193806193806190 | 0.8728771228771220 |
|  |  | 60–90 | 0.9460539460539460 | 0.9460539460539460 |
|  |  | 90–120 | 0.3456543456543450 | 0.8641358641358640 |
|  |  | 120–150 | 0.0079920079920080 | **0.0399600399600399** |
|  | glutamate | -60–0 | 1.0 | — |
|  |  | 0–30 | 0.6983016983016980 | 0.8728771228771220 |
|  |  | 30–60 | 0.2247752247752240 | 0.7492507492507490 |
|  |  | 60–90 | 0.6513486513486510 | 0.8728771228771220 |
|  |  | 90–120 | 0.5284715284715280 | 0.8728771228771220 |
|  |  | **120–150** | 0.0009990009990010 | **0.0099900099900100** |
| **Fall time** | **GABA** | **-60–0** | 1.0 | — |
|  |  | **0–30** | 0.9150849150849150 | 0.9150849150849150 |
|  |  | **30–60** | 0.0789210789210789 | 0.1578421578421570 |
|  |  | **60–90** | 0.0549450549450549 | 0.1373626373626370 |
|  |  | **90–120** | 0.0079920079920080 | **0.0266400266400266** |
|  |  | **120–150** | 0.0009990009990010 | **0.0049950049950050** |
|  | **glutamate** | **-60–0** | 1.0 | — |
|  |  | **0–30** | 0.2287712287712280 | 0.3812853812853810 |
|  |  | **30–60** | 0.5644355644355640 | 0.6271506271506270 |
|  |  | **60–90** | 0.3876123876123870 | 0.4845154845154840 |
|  |  | **90–120** | 0.3426573426573420 | 0.4845154845154840 |
|  |  | **120–150** | — | — |
| **Rise time** | **GABA** | **-60–0** | 1.0 | — |
|  |  | **0–30** | 0.1798201798201790 | 0.1998001998001990 |
|  |  | **30–60** | 0.1088911088911080 | 0.1361138861138860 |
|  |  | **60–90** | 0.0109890109890109 | **0.0156985871271585** |
|  |  | **90–120** | 0.0019980019980020 | **0.0049950049950050** |
|  |  | **120–150** | 0.6643356643356640 | 0.6643356643356640 |
|  | **glutamate** | **-60–0** | 1.0 | — |
|  |  | **0–30** | 0.0109890109890109 | **0.0156985871271585** |
|  |  | **30–60** | 0.0049950049950050 | **0.0099900099900100** |
|  |  | **60–90** | 0.0009990009990010 | **0.0033300033300033** |
|  |  | **90–120** | 0.0009990009990010 | **0.0033300033300033** |
|  |  | **120–150** | 0.0009990009990010 | **0.0033300033300033** |
| **Decay time constant** | **GABA** | **-60–0** | 1.0 | — |
|  |  | **0–30** | 0.8951048951048950 | 0.8951048951048950 |
|  |  | **30–60** | 0.1318681318681310 | 0.3216783216783210 |
|  |  | **60–90** | 0.1608391608391600 | 0.3216783216783210 |
|  |  | **90–120** | 0.0199800199800199 | 0.0666000666000666 |
|  |  | **120–150** | — | — |
|  | **glutamate** | **-60–0** | 1.0 | — |
|  |  | **0–30** | 0.3766233766233760 | 0.5380333951762520 |
|  |  | **30–60** | 0.7402597402597400 | 0.8225108225108220 |
|  |  | **60–90** | 0.3686313686313680 | 0.5380333951762520 |
|  |  | **90–120** | 0.4775224775224770 | 0.5969030969030960 |
|  |  | **120–150** | — | — |
| **Duration (10% peak)** | **GABA** | **-60–0** | 1.0 | — |
|  |  | **0–30** | 0.5714285714285710 | 0.7842157842157840 |
|  |  | **30–60** | 0.6273726273726270 | 0.7842157842157840 |
|  |  | **60–90** | 0.9650349650349650 | 0.9650349650349650 |
|  |  | **90–120** | 0.8731268731268730 | 0.9650349650349650 |
|  |  | **120–150** | — | — |
|  | **glutamate** | **-60–0** | 1.0 | — |
|  |  | **0–30** | 0.0409590409590409 | 0.0999000999000999 |
|  |  | **30–60** | 0.0499500499500499 | 0.0999000999000999 |
|  |  | **60–90** | 0.0239760239760239 | 0.0799200799200799 |
|  |  | **90–120** | 0.0639360639360639 | 0.1065601065601060 |
|  |  | **120–150** | — | — |

**Extended Data Table 13.**
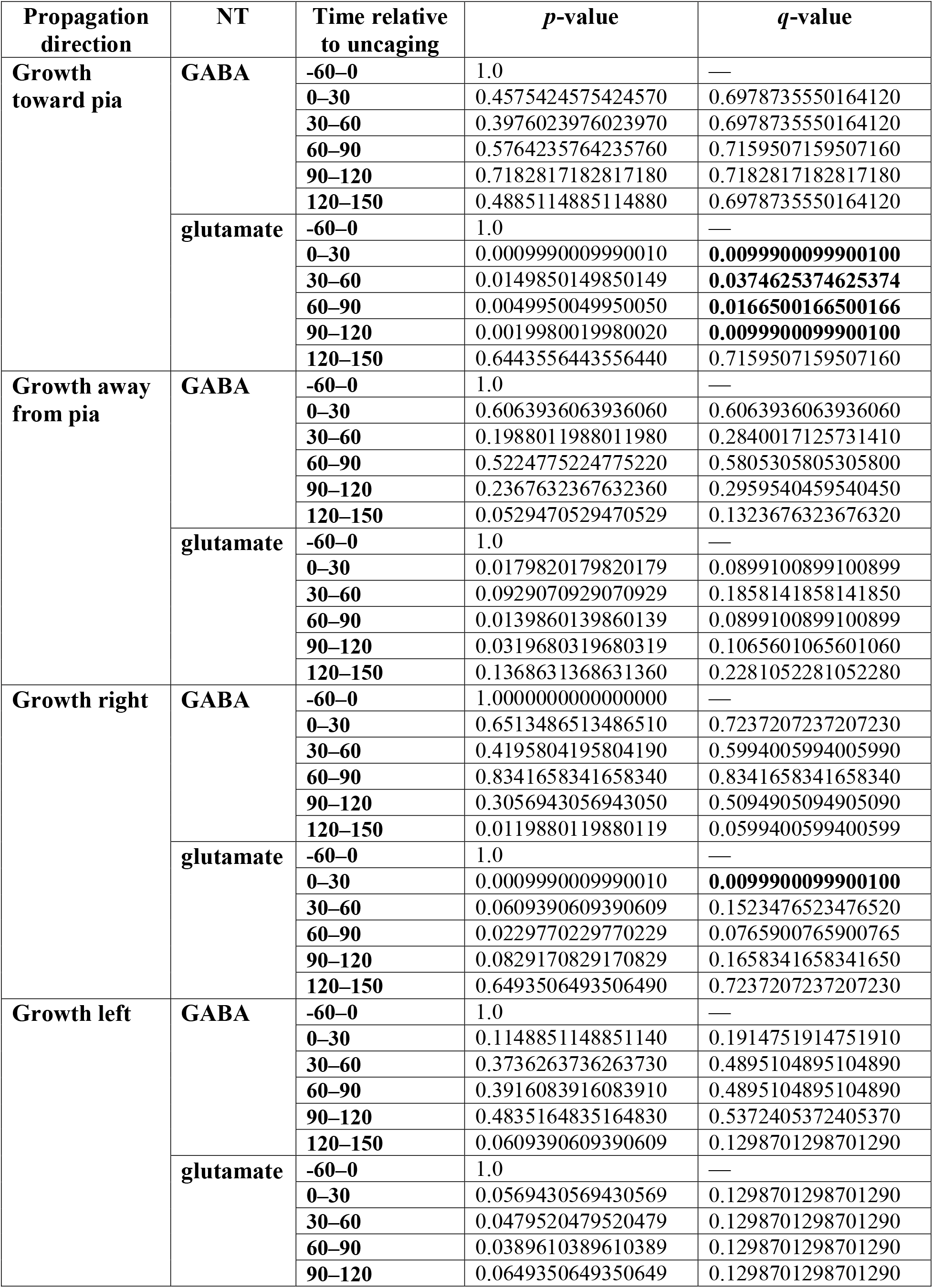
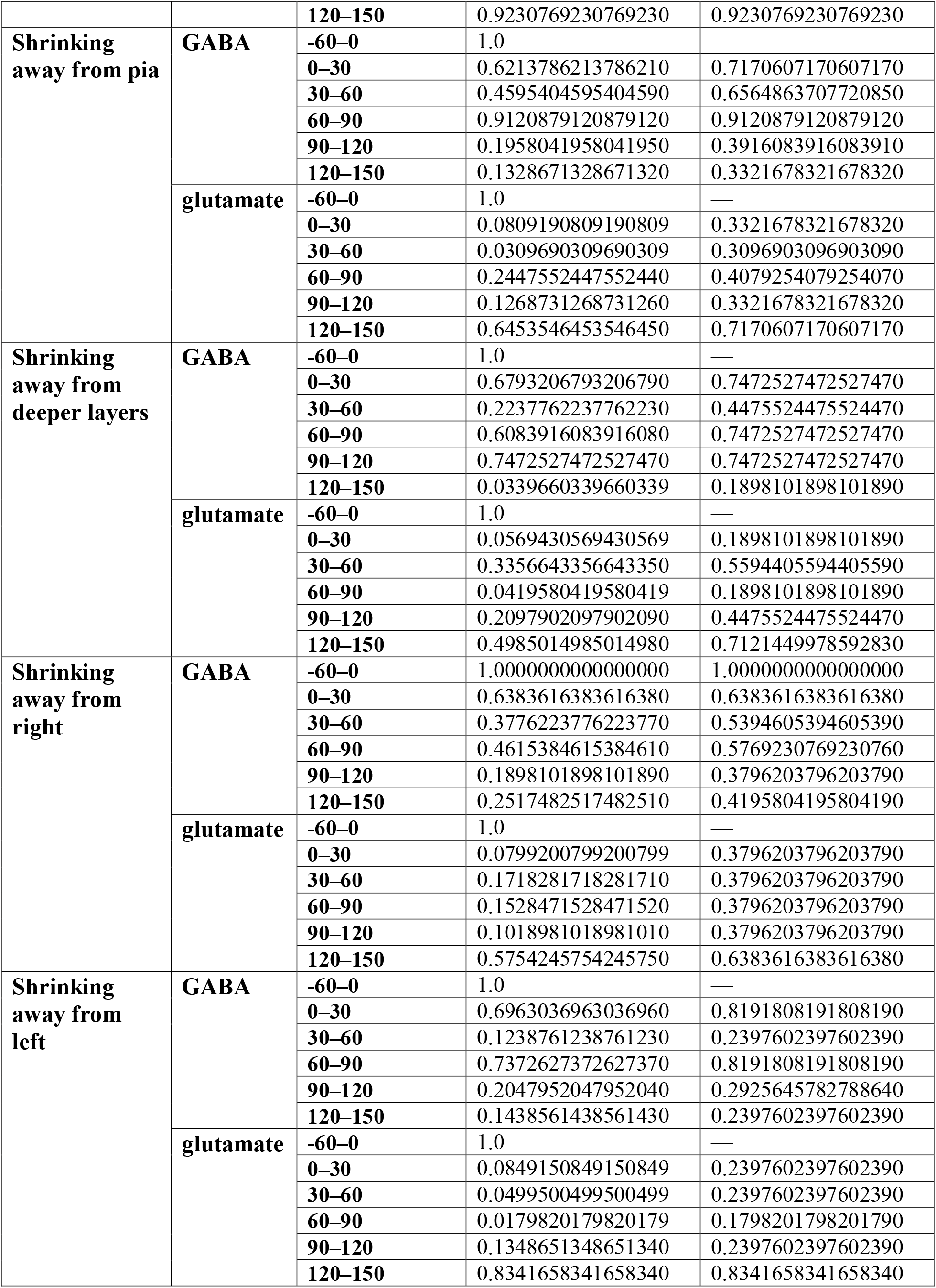
Statistics for Extended Data Fig. 4b: Change in probability of Ca^2+^ events growing or shrinking in the indicated direction compared to baseline. Two-sided permutation testing was used to identify time bins with changes in propagation probability compared to baseline. *p*-values were adjusted across tested time bins using the Benjamini-Hochberg procedure to obtain *q*-values. *q*-values < 0.05 are bold.

**Extended Data Table 14.** Statistics for Extended Data Fig. 7d: Change in event frequency in neighboring cells during multiple rounds of glutamate uncaging or in RuBi-glutamate uncaging controls. Permutation testing used to identify time-points with changes in event frequency greater than chance for each condition. *p*-values corrected for multiple comparisons using Benjamini-Yekutieli procedure with FDR ≤ 0.05. Adjusted *p*-values < 0.05 are bold.

| Dataset | Round | Time relative to uncaging | *p*-values (raw) | *p*-values (adjusted) |
| --- | --- | --- | --- | --- |
| Multi-round glutamate uncaging | Round 1 | 0–30 | 0.107389261073893 | 0.482586244333393 |
|  |  | 30–60 | 0.497850214978502 | 1.86436755938545 |
|  |  | 60–90 | 0.373962603739626 | 1.48280689599646 |
|  |  | 90–120 | 0.101789821017898 | 0.482586244333393 |
|  |  | 120–150 | 0.613638636136386 | 2.17703012457662 |
|  |  | 150–180 | 0.00779922007799220 | **0.0438102037453499** |
|  |  | 180–210 | 0.0619938006199380 | 0.321447648782449 |
|  |  | 210–240 | 0.00389961003899610 | **0.0238964747701909** |
|  |  | 240–270 | 0.135986401359864 | 0.572902664362268 |
|  |  | 270–300 | 0.00119988001199880 | **0.00898670846058460** |
|  |  | 300–330 | 0.000199980001999800 | **0.00192572324155384** |
|  |  | 330–360 | 0.000799920007999200 | **0.00674003134543845** |
|  |  | 360–390 | 0.00289971002899710 | **0.0195460909017715** |
|  |  | 390–420 | 9.99900009999000e-05 | **0.00112333855757308** |
|  |  | 420–450 | 9.99900009999000e-05 | **0.00112333855757308** |
|  |  | 450–480 | 9.99900009999000e-05 | **0.00112333855757308** |
|  |  | 480–510 | 9.99900009999000e-05 | **0.00112333855757308** |
|  |  | 510–540 | 9.99900009999000e-05 | **0.00112333855757308** |
|  |  | 540–570 | 9.99900009999000e-05 | **0.00112333855757308** |
|  | Round 2 | 0–30 | 0.104989501049895 | 0.505502350907884 |
|  |  | 30–60 | 0.379962003799620 | 1.58866503536305 |
|  |  | 60–90 | 0.679432056794321 | 2.41044805222391 |
|  |  | 90–120 | 0.561243875612439 | 2.10176644121922 |
|  |  | 120–150 | 0.363463653634637 | 1.58866503536305 |
|  |  | 150–180 | 0.0125987401259874 | 0.0943604388361383 |
|  |  | 180–210 | 0.0209979002099790 | 0.141540658254208 |
|  |  | 210–240 | 0.400659934006599 | 1.58866503536305 |
|  |  | 240–270 | 0.0349965003499650 | 0.214455542809405 |
|  |  | 270–300 | 0.103689631036896 | 0.505502350907884 |
|  |  | 300–330 | 0.0536946305369463 | 0.301616402708371 |
|  |  | 330–360 | 0.000199980001999800 | **0.00168500783635961** |
|  |  | 360–390 | 9.99900009999000e-05 | **0.000962861620776922** |
|  |  | 390–420 | 9.99900009999000e-05 | **0.000962861620776922** |
|  |  | 420–450 | 9.99900009999000e-05 | **0.000962861620776922** |
|  |  | 450–480 | 9.99900009999000e-05 | **0.000962861620776922** |
|  |  | 480–510 | 9.99900009999000e-05 | **0.000962861620776922** |
|  |  | 510–540 | 9.99900009999000e-05 | **0.000962861620776922** |
|  |  | 540–570 | 9.99900009999000e-05 | **0.000962861620776922** |
|  | Round 3 | 0–30 | 0.684231576842316 | 2.71306085275502 |
|  |  | 30–60 | 0.654234576542346 | 2.71306085275502 |
|  |  | 60–90 | 0.806319368063194 | 3.01953404275643 |
|  |  | 90–120 | 0.357364263573643 | 1.85299015604593 |
|  |  | 120–150 | 0.244075592440756 | 1.49567422856502 |
|  |  | 150–180 | 0.354064593540646 | 1.85299015604593 |
|  |  | 180–210 | 0.622237776222378 | 2.71306085275502 |
|  |  | 210–240 | 0.904009599040096 | 3.20719070495311 |
|  |  | 240–270 | 0.0222977702229777 | 0.187878373754097 |
|  |  | 270–300 | 0.479252074792521 | 2.30749787419189 |
|  |  | 300–330 | 0.135886411358864 | 1.01774473316121 |
|  |  | 330–360 | 0.210678932106789 | 1.42012460448388 |
|  |  | 360–390 | 0.00909909009099090 | 0.0876204074906999 |
|  |  | 390–420 | 0.000599940005999400 | **0.00674003134543845** |
|  |  | 420–450 | 9.99900009999000e-05 | **0.00168500783635961** |
|  |  | 450–480 | 9.99900009999000e-05 | **0.00168500783635961** |
|  |  | 480–510 | 0.000499950004999500 | **0.00674003134543845** |
|  |  | 510–540 | 9.99900009999000e-05 | **0.00168500783635961** |
|  |  | 540–570 | 9.99900009999000e-05 | **0.00168500783635961** |
| RuBi-glutamate uncaging control | Round 1 | 0–30 | 0.982401759824018 | 3.53816171786332 |
|  |  | 30–60 | 0.778222177782222 | 3.08574493891456 |
|  |  | 60–90 | 0.0261973802619738 | 0.346630183479692 |
|  |  | 90–120 | 0.00819918008199180 | 0.276341285162977 |
|  |  | 120–150 | 0.392460753924608 | 1.83778188018955 |
|  |  | 150–180 | 0.997300269973003 | 3.53816171786332 |
|  |  | 180–210 | 0.408959104089591 | 1.83778188018955 |
|  |  | 210–240 | 0.601639836016398 | 2.53467303784395 |
|  |  | 240–270 | 0.0212978702129787 | 0.346630183479692 |
|  |  | 270–300 | 0.211778822117788 | 1.18961553246989 |
|  |  | 300–330 | 0.103589641035896 | 0.634788406715840 |
|  |  | 330–360 | 0.0359964003599640 | 0.346630183479692 |
|  |  | 360–390 | 0.0818918108189181 | 0.552008567191409 |
|  |  | 390–420 | 0.0743925607439256 | 0.552008567191409 |
|  |  | 420–450 | 0.00609939006099390 | 0.276341285162977 |
|  |  | 450–480 | 0.0521947805219478 | 0.439787045289859 |
|  |  | 480–510 | 0.0229977002299770 | 0.346630183479692 |
|  |  | 510–540 | 0.0346965303469653 | 0.346630183479692 |
|  |  | 540–570 | 0.267973202679732 | 1.38948338505962 |
|  | Round 2 | 0–30 | 0.0592940705929407 | 1.75240814981400 |
|  |  | 30–60 | 0.604039596040396 | 2.85187576303865 |
|  |  | 60–90 | 0.0238976102389761 | 1.61086749155979 |
|  |  | 90–120 | 0.121987801219878 | 1.75240814981400 |
|  |  | 120–150 | 0.916208379162084 | 3.25046880096066 |
|  |  | 150–180 | 0.847915208479152 | 3.17530365607323 |
|  |  | 180–210 | 0.755224477552245 | 2.99455627953510 |
|  |  | 210–240 | 0.137786221377862 | 1.75240814981400 |
|  |  | 240–270 | 0.155984401559844 | 1.75240814981400 |
|  |  | 270–300 | 0.676932306769323 | 2.85187576303865 |
|  |  | 300–330 | 0.207279272072793 | 1.99601213987056 |
|  |  | 330–360 | 0.479252074792521 | 2.69208085322388 |
|  |  | 360–390 | 0.116288371162884 | 1.75240814981400 |
|  |  | 390–420 | 0.341065893410659 | 2.55447187992117 |
|  |  | 420–450 | 0.324367563243676 | 2.55447187992117 |
|  |  | 450–480 | 0.666833316668333 | 2.85187576303865 |
|  |  | 480–510 | 0.419858014198580 | 2.57285378359055 |
|  |  | 510–540 | 0.604739526047395 | 2.85187576303865 |
|  |  | 540–570 | 0.381361863813619 | 2.57064795515023 |
|  | Round 3 | 0–30 | 0.593640635936406 | 2.66770440652454 |
|  |  | 30–60 | 0.869013098690131 | 3.36402453374550 |
|  |  | 60–90 | 0.287671232876712 | 2.66770440652454 |
|  |  | 90–120 | 0.0991900809919008 | 1.67152777366874 |
|  |  | 120–150 | 0.327267273272673 | 2.66770440652454 |
|  |  | 150–180 | 0.999100089991001 | 3.54454701071690 |
|  |  | 180–210 | 0.505849415058494 | 2.66770440652454 |
|  |  | 210–240 | 0.406459354064594 | 2.66770440652454 |
|  |  | 240–270 | 0.0353964603539646 | 1.19298554814261 |
|  |  | 270–300 | 0.515848415158484 | 2.66770440652454 |
|  |  | 300–330 | 0.357564243575642 | 2.66770440652454 |
|  |  | 330–360 | 0.0989901009899010 | 1.67152777366874 |
|  |  | 360–390 | 0.572942705729427 | 2.66770440652454 |
|  |  | 390–420 | 0.465453454654535 | 2.66770440652454 |
|  |  | 420–450 | 0.859514048595141 | 3.36402453374550 |
|  |  | 450–480 | 0.206879312068793 | 2.66770440652454 |
|  |  | 480–510 | 0.00809919008099190 | 0.545942538980515 |
|  |  | 510–540 | 0.431456854314569 | 2.66770440652454 |
|  |  | 540–570 | 0.898310168983102 | 3.36402453374550 |

